# Bilateral integration in somatosensory cortex is controlled by behavioral relevance

**DOI:** 10.1101/2024.05.28.596311

**Authors:** Hyein Park, Hayagreev V.S. Keri, Chaeyoung Yoo, Chengyu Bi, Scott R. Pluta

**Affiliations:** Purdue University, Department of Biological Sciences, West Lafayette, IN, 47907

## Abstract

Sensory perception naturally requires processing stimuli from both sides of the body. Yet, how neurons bind stimulus features across the hemispheres to create a unified perceptual experience remains unknown. To address this question, we performed large-scale recordings from neurons in both somatosensory cortices (S1) while mice shared information between their hemispheres and discriminated between two categories of bilateral stimuli. When expert mice touched stimuli associated with reward, they moved their whiskers with greater bilateral symmetry. During this period, synchronous spiking and enhanced spike-field coupling emerged between the hemispheres. This coordinated activity was absent in stimulus-matched naïve animals, indicating that interhemispheric (IH) binding was controlled by a goal-directed, internal process. In S1 neurons, the addition of ipsilateral touch primarily facilitated the contralateral, principal whisker response. This facilitation primarily emerged for reward-associated stimuli and was lost on trials where expert mice failed to respond. Taken together, these results reveal a novel state-dependent logic underlying bilateral integration in S1, where stimulus binding and facilitation are controlled by behavioral relevance.

## Introduction

Tactile perception naturally involves coordinated movements guided by sensory feedback on both sides of the body ^1–4^. For instance, humans use both of their hands to interact with objects, and rodents use whiskers on both sides of their face to navigate complex terrain. This bilateral coordination depends on the ability to bind stimuli across the hemispheres and create a unified perceptual experience^5^. However, the neural computations and brain areas supporting bilateral perception are unclear. Few studies have investigated bilateral integration in primary somatosensory cortex (S1), where touch is integrated from both sides of the body for the first time ^6–8^. Studies in anesthetized and non-behaving animals have revealed the presence of ipsilateral receptive fields in S1 ^9–11^, yet the paucity of ipsilateral-evoked spiking has led to the hypothesis that bilateral integration must occur in areas upstream ^12^. However, this hypothesis is challenged by deficits in bilateral coordination caused by severing the parietal section of the corpus callosum ^1,13–16^. Given the delays imposed by sequential processing, combined with the speed of tactile sensation, we hypothesized that bilateral integration must emerge in S1 neurons during behaviors that require bilateral coordination ^17^.

Multiple studies have demonstrated that sensory associations and experience shape ipsilateral responses in S1 ^18,19^. For instance, ipsilateral touch responses increase after reinforcement learning and during stimulus anticipation, suggesting a state-dependent change in the effectiveness of callosal inputs ^20,21^. Repetitive stimulation of homotopic neurons in the opposite somatosensory cortex enhances ipsilateral touch responses, indicating a purely activity-driven strengthening of callosal input ^22^. Overall, multiple forms of experience-dependent IH plasticity have been observed ^23–29^, implying that information flow between S1s is inherently flexible.

In anesthetized and non-behaving animals, stimulus competition is the default mode of bilateral integration. Under anesthesia, ipsilateral stimuli largely suppress the contralateral response. In awake quiescent animals, the probability of observing bilateral facilitation increases, yet remains less prevalent than suppression ^6,17,30,31^. Similarly, optogenetic stimulation of S1 callosal axons largely inhibits contralateral touch responses ^32,33^. Despite these competitive interactions, sensory associations learned in one hemisphere are transmitted through the callosum and more quickly learned in the opposite hemisphere ^13,34^. Recent evidence reveals that S1 callosal circuits increase resting-state correlations between the hemispheres, providing further support for IH communication during unstimulated periods ^35^. Given these results, the prevailing notion is that callosal circuits in S1 enforce bilateral stimulus competition while paradoxically having the capacity to transfer sensory associations ^8,13^. However, this assessment is insufficient, since tactile behaviors require bilateral stimulus coordination in real-time ^1,2,5,36–38^. Given the state-dependent changes to ipsilateral touch responses in S1, we hypothesized that S1 neurons switch from performing bilateral stimulus competition (no interaction or suppression) to bilateral stimulus coordination (facilitation and binding) in accordance with behavioral relevance.

To reveal the state-dependent logic of bilateral integration in S1, we developed a novel task where mice used active whisker touch to share information between their hemispheres and discriminate between two categories of bilateral stimuli. We made large-scale multi-site recordings from neurons in both somatosensory cortices while tracking animal movement and choice. We also recorded naïve mice that performed the same active bilateral touches without any sensory associations. First, we discovered that bilateral whisking symmetry was enhanced in expert mice by the expectation of reward. In S1 neurons, bilateral facilitation emerged during reward-associated bilateral touch, while in naïve mice, most neurons lacked any significant bilateral interaction. In expert mice, spike timing became highly synchronous across the hemispheres and was phase-locked to the field potential in the opposite hemisphere, indicating that IH communication was controlled by an internal, goal-directed process. Bilateral integration was lost on trials when expert mice failed to report the stimulus, revealing the necessity of task engagement. Taken together, these data indicate that the somatosensory cortices switch from two lateralized spatial representations in a naïve state to a single unified representation during bilateral coordination.

## Results

### Bilateral whisking symmetry reflects reward expectation and behavioral context

To reveal the neural mechanisms underlying bilateral integration in S1, we developed a Go/No-Go task where mice were rewarded for discriminating the spatial relationship between two objects, with one object presented to each side of their face (Fig. 1 and Suppl. Fig. 1). Mice solved the task using active touch and reported the presence of a Go stimulus by licking a water port (Fig. 1a). Before training began, the Go stimuli for each mouse was designated (and remained) as either homotopic (HM, matching) or heterotopic (HT, nonmatching) whisker pairs (Fig. 1b). Since HM and HT stimuli (four total) were presented across both bilateral pairs of whiskers (Fig. 1b), mice could not solve the task using a single whisker, thereby requiring bilateral coordination. Mice initiated a trial by locomoting on a circular treadmill for a short distance (∼110 cm, Fig. 1a, c & Suppl. Video 1). Stimuli were presented by pneumatically extending the touch surfaces into the anterior region of their whisking field with single-whisker accuracy^39^. The experiment took place in a sound attenuated and dark enclosure containing white noise. While mice performed the task, we made bilateral, high-density electrophysiological recordings from their primary somatosensory cortices (S1) and imaged each whisker pad at 500 frames per second under infrared illumination. Using intrinsic imaging and multi-site silicon probes, we targeted our recordings to all four of the S1 barrel columns directly involved in the task (C1 and D1 on each side, Suppl. Fig. 1i). We successfully recorded from ten expert mice, with six mice designated as HM-Go and four designated as HT-Go (159 ± 11 neurons per mouse). Removing the ability of mice to touch the surfaces, via whisker trimming, reduced performance to chance, demonstrating the necessity of active touch to solve the task (Fig. 1d and Suppl. Fig. 1c). To further examine the role of behavioral relevance on bilateral processing, we made additional recordings from seven naïve mice that performed the same active touches but without sensory associations (121 ± 30 neurons per mouse). We also recorded four mice that learned an association between active touch and reward but were not performing bilateral discrimination (62 ± 17 touch-responsive neurons per mouse).

**Figure 1.**
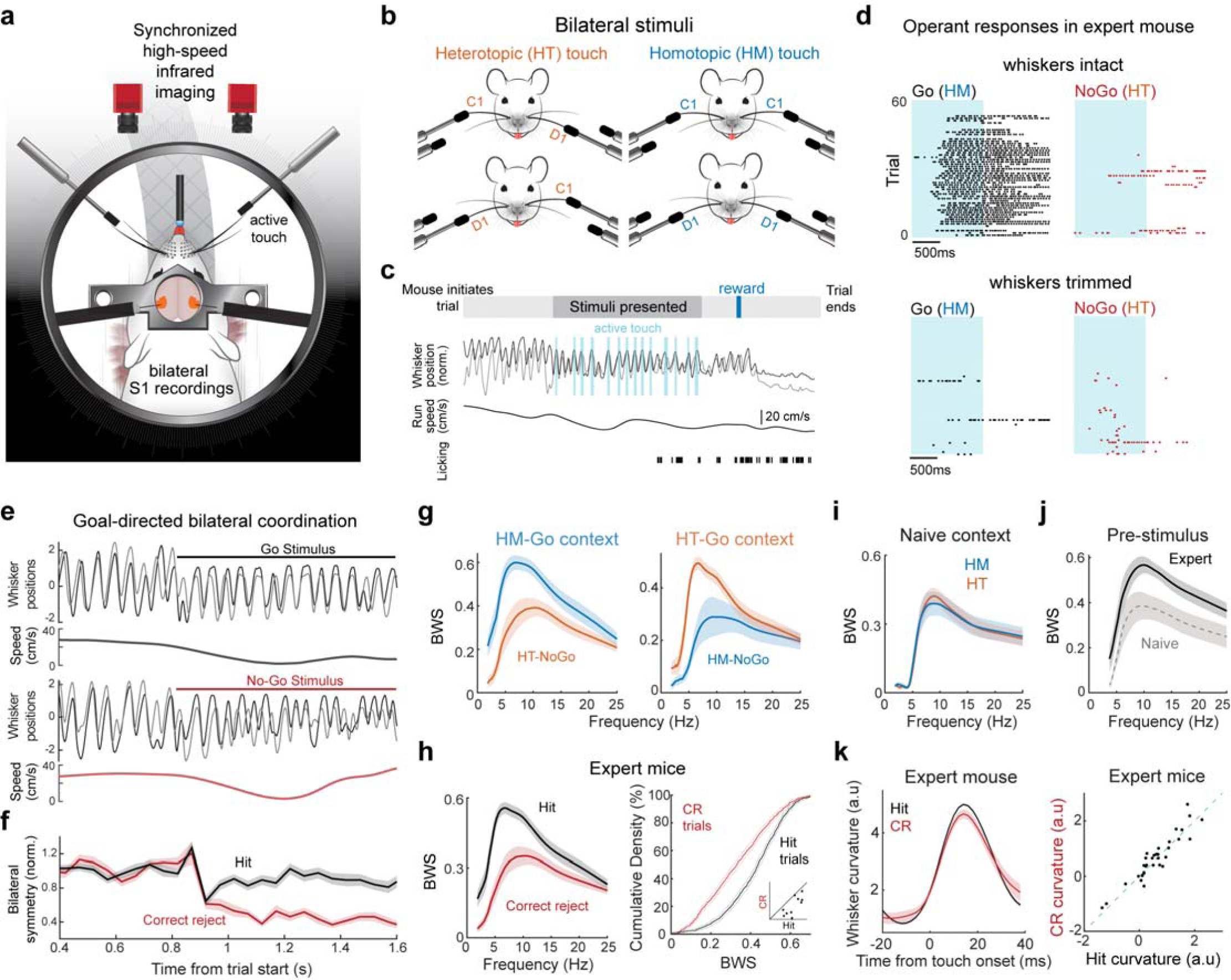
Bilateral whisking symmetry reflects task engagement and reward expectation. (**a**) Schematic of the recording setup. Single-unit activity from S1 neurons in both hemispheres were recorded while mice performed bilateral discrimination using active touch. Infrared high-speed (500 fps) cameras imaged bilateral whisker motion and touch dynamics. (**b**) Illustrations of the four different bilateral stimuli. Left, heterotopic (HT) touch: different whiskers on each side of a mouse’s face are stimulated (C1-D1 or D1-C1 pairs). Right, homotopic (HM) touch: same whiskers on both sides are stimulated (C1-C1 or D1-D1 pairs). (**c**) Trial structure (top) and traces of whisking, touch times, locomotion, and licking during an example trial. Stimulus presentation lasted 1350 ms. (**d**) Example lick rasters from an expert mouse with C1 & D1 whiskers intact (top), and with whiskers trimmed off near their base (bottom). Blue shadow indicates stimulus presence. (**e**) Example whisking and locomotion traces during a Go (top) and a No-Go (bottom) trial. (**f**) Average bilateral symmetry during whisking from the mouse in **e** during hit (black) and correct rejection (CR, red) trials. (**g**) Bilateral whisking symmetry (BWS) calculated during stimulus presentation for all expert HM-Go mice (p = 5.5e^−52^, two-sample t-test; 552 (46×12) samples per condition from 46 frequency points across 12 whisker pairs in 6 mice) and HT-Go mice (p = 1.9e^−23^, two-sample t-test; 368 samples per condition across 8 whisker pairs in 4 mice). (**h**) Bottom: Left, BWS of all expert mice during hit and CR trials (p = 3.4e^−68^, two-sample t-test; 920 samples for hit and CR respectively, 20 whisker pairs in 10 mice). Right, cumulative density of BWS per trial in all expert mice. Right bottom shows average BWS per mouse. (**i**) BWS of naïve mice during HM and HT touch (p = 0.9747, two-sample t-test; 644 samples per condition across 14 whisker pairs in 7 mice). (**j**) BWS of all expert (black, 10 mice) and naïve mice (gray, 7 mice) during the pre-stimulus period (p = 1.3e^−18^, two-sided Wilcoxon rank-sum test; 340 samples from 10 expert mice; 238 samples from 7 naïve mice). (**k**) Left, average whisker curvature during touch in an example mouse. Right, scatter plot of average whisker curvature for individual whiskers (p = 0.13, Wilcoxon signed rank test, 10 mice, 40 whiskers). All error bars represent the mean ± s.e.m.

Since task-specific adjustments to active sensing are a hallmark of skilled sensorimotor learning^40^, we first examined whether the whisking strategy of mice reflected the behavioral relevance of the stimulus. We reasoned that if mice were solving the task by binding stimulus features across their hemispheres, they would adopt a whisking strategy that maximizes bilateral coordination. Interestingly, we discovered that bilateral whisking displayed significantly greater symmetry for Go (Hit) compared to No-Go (CR, correct rejection) stimulus periods, across both HT- and HM-Go contexts (Fig. 1e, f). To better understand this movement symmetry across stimulus conditions and behavioral contexts, we employed a metric we call ‘bilateral whisking symmetry’ (BWS), that was calculated from the angular phase consistency between HT and HM whisker pairs ^41,42^. BWS on CR trials was significantly lower than Hit trials (Fig. 1g, h), revealing that mice deliberately reduced their bilateral coordination after deciding the stimulus was not associated with reward. In agreement with this finding, BWS during CR touch periods was similar to BWS in naïve animals, which had no associative training (Fig. 1i).

Furthermore, expert mice displayed significantly greater BWS than naïve mice during trial initiation (pre-stimulus, Fig. 1j), suggesting that movement symmetry was a deliberate strategy for bilateral discrimination. To determine if BWS was motivated by the bilateral context of our task, or if was simply a reflection of reward expectation, we investigated mice that associated touch with reward, but were not performing bilateral discrimination, and found they had significantly lower BWS (Suppl. Fig. 1f, g). Despite these goal-directed changes in movement, the strength of active touch (as indicated by whisker curvature) was equivalent across conditions (Fig. 1k). Taken together, these data suggest that BWS is driven by both reward expectation and active engagement in bilateral discrimination. Are these internally motivated changes to active sensing reflected in the bilateral integration of S1 neurons?

### Bilateral facilitation in S1 neurons is controlled by behavioral relevance

To determine the impact of behavioral relevance on bilateral integration in S1 neurons, we aligned spiking to the onset of active touch. This approach enabled us to relate neuronal firing rates to a fundamental stimulus feature. To identify the principal whisker (PW) of each neuron, mice were periodically presented single-whisker stimuli, which had no impact on task performance (Fig. 2a). We discovered that the PW response in S1 neurons was enhanced by the addition of ipsilateral touch, but only when the bilateral stimulus was associated with reward (Fig. 2b, c). To illustrate this effect across the population, we calculated a bilateral integration index, based on the normalized difference between the PW and bilateral responses (see methods). In expert mice, S1 neurons had significantly greater bilateral facilitation for their respective Go (Hit) than No-Go (CR) stimuli (Fig. 2c and Fig. 2d, left). In naïve mice, the S1 population did not show significant bilateral integration, regardless of stimulus configuration (Fig. 2d, right). The probability of an individual neuron displaying significant bilateral facilitation doubled in expert mice (Fig. 2e), and 78% of S1 neurons in expert mice preferred bilateral over PW touch (Fig. 2f). Importantly, the S1 population in mice that were expecting reward, but not performing bilateral discrimination, had no significant bilateral integration (Suppl. Fig. 2e). Taken together, our data from naïve mice are consistent with previous reports in non-behaving animals, while our data in bilaterally discriminating mice reveal that bilateral integration in S1 is controlled by behavioral relevance. These goal-directed gains in bilateral integration could not be explained by differences in the interhemispheric touch interval (ITI) or the total number of touches in the trial (Suppl. Fig. 2a, b). However, bilateral facilitation decreased when ipsilateral touch preceded contralateral touch by more than 12 ms, indicating a finite window of ipsilateral-mediated facilitation.

**Figure 2.**
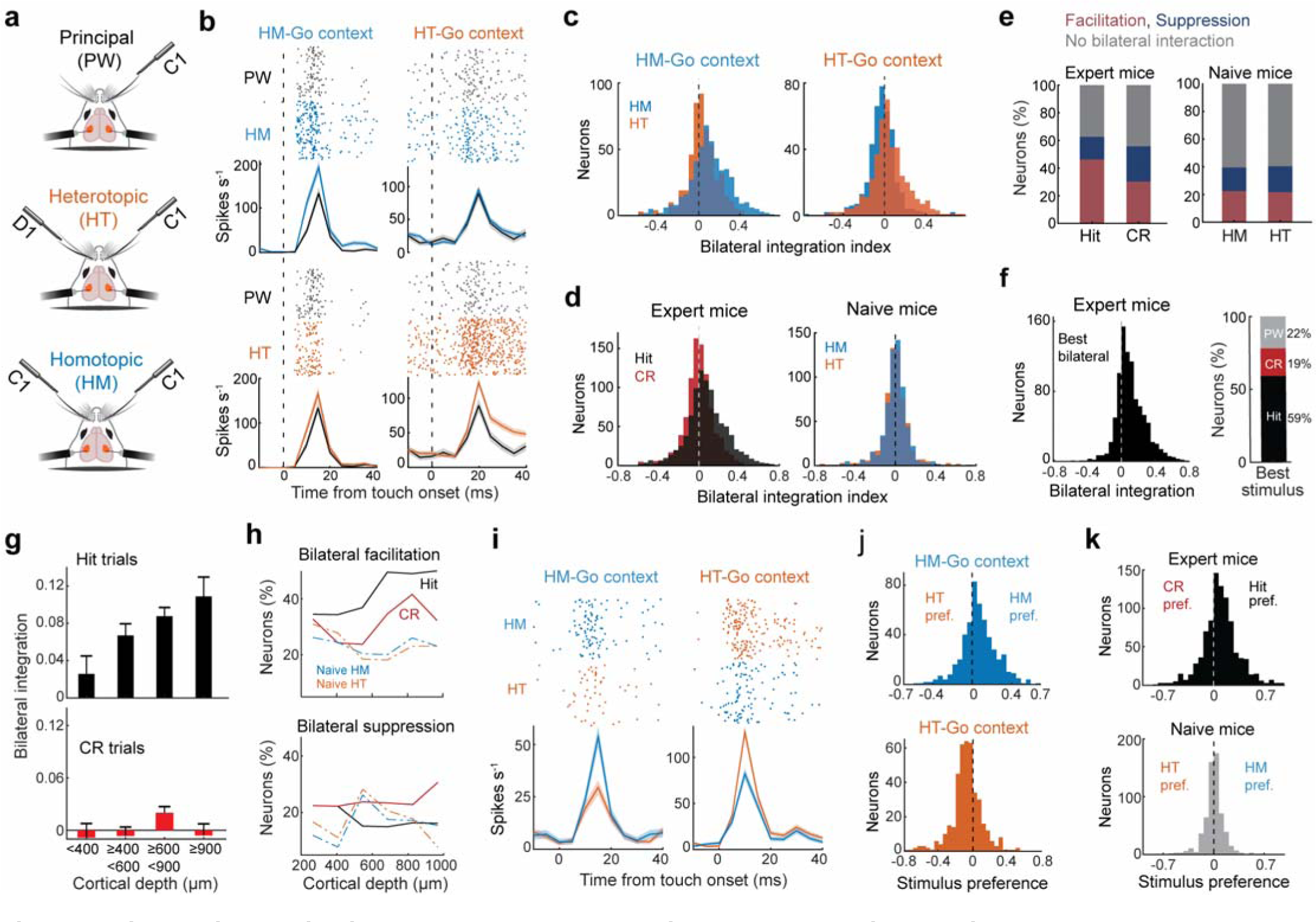
Bilateral integration in S1 neurons emerges during reward-associated active touch. (**a**) Schematic illustrating an example comparison between unilateral and bilateral touch conditions. Responses to principal whisker (PW) touch were compared with corresponding heterotopic (HT) and homotopic (HM) bilateral touches. (**b**) Example neurons from an expert HM-go mouse (left column) and an expert HT-go mouse (right column), comparing responses to PW and bilateral touches. (**c**) Distributions of bilateral integration indices for the HM-go and HT-go behavioral contexts; Neurons from expert HM-go mice (left, p = 1.8e^−15^, paired-sample t-test; 6 mice, 503 neurons); Neurons from expert HT-go mice (right, p = 8.0e^−16^ paired-sample t-test; 4 mice, 411 neurons) (**d**) Distributions of bilateral integration indices for the Hit and CR conditions in all expert mice (p = 1.1e^−29^, paired-sample t-test; 10 mice, 914 neurons), and for the HM and HT conditions in naïve mice (right, p = 0.64, paired-sample t-test; 7 mice, 609 neurons). The CR population distribution was not significantly different from zero (p = 0.2, one-sample t-test; Left in red) (**e**) Percentage of neurons with significant bilateral facilitation or suppression (10 expert mice, 914 neurons; ANOVA with Tukey comparison between PW, HM, and HT responses, α = 0.05). In naïve mice, differences between PW, HM and HT touch responses were also tested for significance (7 naïve mice, 609 neurons; ANOVA with Tukey comparison, α = 0.05). (**f**) Left: Bilateral integration index using best bilateral stimulus, defined as a stimulus resulting in the highest bilateral integration index for a given neuron. Right: Percentage of neurons categorized as Hit-, CR-, and PW-best. (**g**) Bilateral integration indices of neurons in expert mice across cortical depth (10 mice, 914 neurons) during hit trials (p = 0.013, one-way ANOVA; top) and CR trials (p = 0.08, one-way ANOVA; right). (**h**) Percentage of neurons that show significant bilateral facilitation (top) and suppression (bottom) across cortical depth (expert, 10 mice, 914 neurons; naïve, 7 mice, 609 neurons). (**i**) Two example neurons during HM and HT touch. Left: Neuron from an HM-go mouse. Right: Neuron from an HT-go mouse. (**j**) Distributions of bilateral stimulus preference (HM vs. HT) in neurons from HM-Go (p = 2e^−15^, one-sample t-test; 6 mice, 503 neurons) and HT-Go mice (p = 1.2e^−15^, one-sample t-test; 4 mice, 411 neurons). (**k**) Distributions of bilateral stimulus preference in all expert mice (top, p = 2.2e^−29^, one-sample t-test; 10 mice, 914 neurons) and all naïve mice (p = 0.8, one-sample t-test; 7 mice, 609 neurons). All error bars represent the mean ± s.e.m.

Since our linear probes spanned 840 µm of vertical space, we next examined bilateral integration as a function of cortical depth. We discovered that bilateral integration progressively increased with depth due to an increasing likelihood of bilateral facilitation and a decreasing likelihood of bilateral suppression, but only for stimuli associated with reward (Fig. 2g, h and Suppl. Fig. 3). Bilateral integration was similar between putative pyramidal cells and putative fast-spiking interneurons (Suppl. Fig. 4).

While the above data demonstrate that the impact of the ipsilateral stimulus depends on learned associations, our task did not require mice to distinguish unilateral (PW) from bilateral stimuli. To determine how well S1 neurons differentiated between the task-relevant stimuli, we next compared the HM and HT touch responses across the different behavioral contexts (Fig. 2g). S1 neurons preferred the bilateral stimulus that was associated with reward, when comparing HM and HT stimuli with the same contralateral whisker (Fig. 2i, j, k top), while S1 neurons in naïve mice showed no stimulus preference (Fig. 2k, bottom). To determine if learning the task altered ipsilateral integration in S1 neurons, we calculated the probability of observing an ipsilateral touch response in expert and naïve mice. S1 neurons in expert mice were nearly twice as likely as naïve mice to significantly respond to unilateral ipsilateral touch (9 ± 2.4% in expert vs. 5 ± 1.5% in naïve; Suppl. Fig. 2f – h).

### Interhemispheric spike synchrony is enhanced during bilateral discrimination

The neural mechanisms of perception extend beyond the firing rate of individual neurons. Growing evidence suggests that precise patterns of coordinated spikes distributed across distant populations of neurons are essential for encoding stimulus features and optimally driving downstream decision centers ^43–45^. To determine the role of coordinated spiking in bilateral perception, we tested the impact of behavioral relevance on spike synchrony between the somatosensory cortices. To do so, we first created a spike-triggered average of IH spike intervals between pairs of S1 neurons in opposite hemispheres that were significantly driven by the stimulus (Fig. 3a). For each pair of neurons, a jittered (±75 ms) distribution of IH spike intervals was subtracted from the real distribution (see Methods), to remove the effect of neuronal firing rates. We discovered that IH spike intervals were much more likely to be near-zero when mice correctly sensed the Go stimulus, for both HM-Go and HT-Go behavioral contexts. We expressed these IH spike intervals in terms of a fractional change in IH firing rate (4 ms bin size), by first subtracting off and then dividing by the jittered distribution (Fig. 3b). Across the population of S1 neurons, IH firing rates were significantly greater on Hit than CR trials (Fig. 3C).

**Figure 3.**
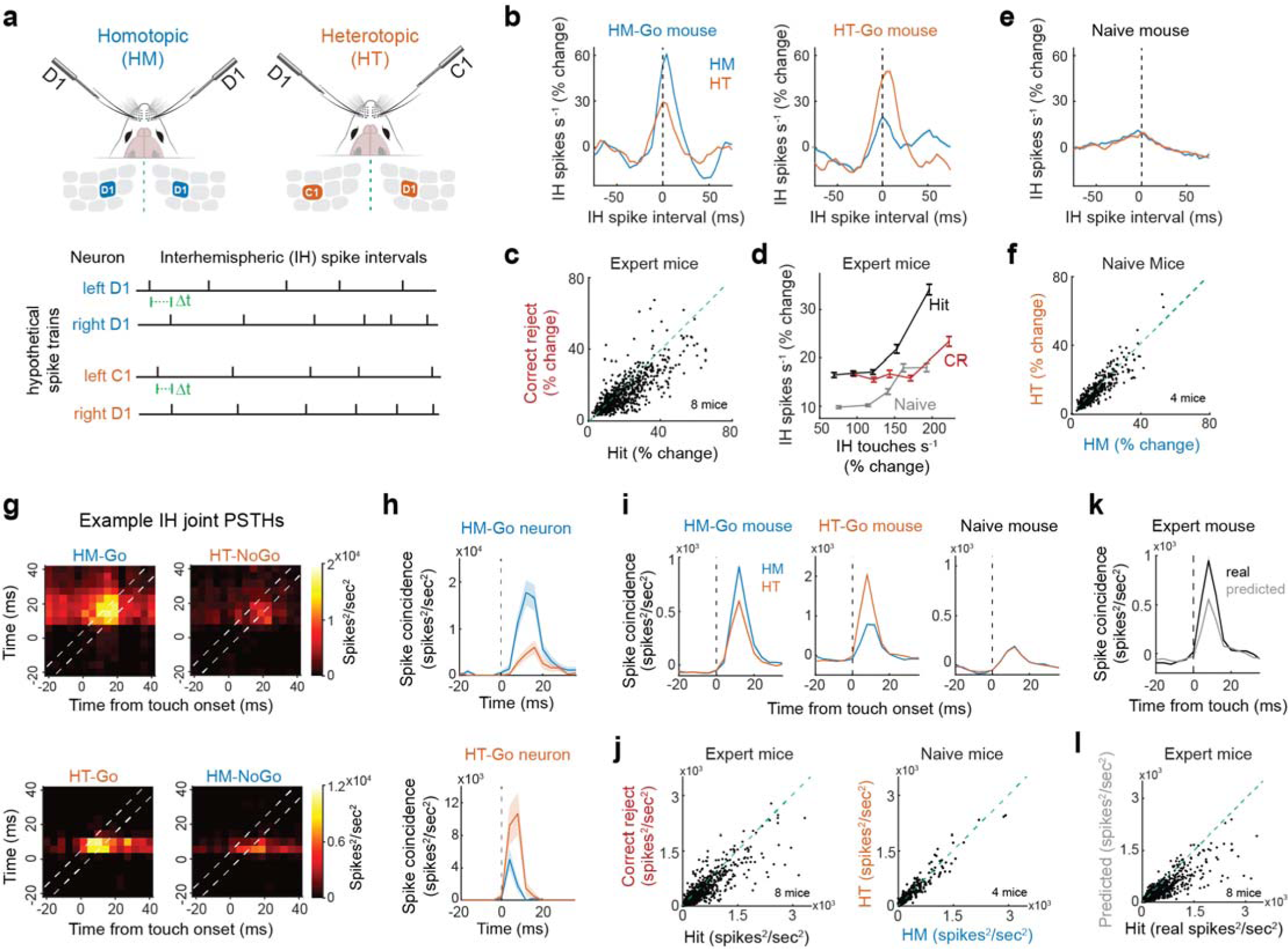
Interhemispheric spike synchrony is controlled by behavioral relevance. (**a**) Schematic illustrating the calculation of interhemispheric time lags between pairs of S1 neuronal spike trains during HM and HT touch. (**b**) Population averaged interhemispheric firing rates as a function of temporal lag between pairs of neurons. Left, example HM-Go mouse during HM (4142 neuron pairs) and HT (3315 neuron pairs) touches. Right, example HT-Go mouse during HM (909 neuron pairs) and HT (1129 neuron pairs) touches. (**c**) Scatter plot comparing population-level IH firing rates during Hit and CR trials in expert mice (p = 5e^−40^, signed-rank test, 8 mice, 783 neurons). (**d**) The relationship between IH touch rate and IH spike rate during Hit and CR trials in expert mice and bilateral trials in naïve mice. (**e**) IH firing rate during HM (blue, 1953 neuron pairs) and HT (orange, 1989 neuron pairs) touch in naïve mice. (**f**) Scatter plot comparing population-level IH firing rates during HM (blue) and HT (orange) touch in naive mice (p = 0.83, signed-rank test, 4 mice, 409 cells). (**g**) Heatmap of spike coincidence (spikes^2^/sec^2^) between two neurons in opposite hemispheres during HM-Go (top) and HT-Go (bottom) behavioral contexts. Only IH touch intervals ≤10 ms were used. (**h**) Joint-PSTH taken from the diagonal of **g** during HM and HT touch conditions. (**i**) Joint-PSTH of average spike coincidence in an example HM-Go (3469 neuron pairs during HM and 3393 neuron pairs during HT), HT-Go (909 neuron pairs during HM and 1129 neuron pairs during HT) and naïve mouse (2081 neuron pairs during HM and 2098 neuron pairs during HT). (**j**) Left, population-level scatter plot comparing spike coincidence between Hit (black) and CR (red) trials in expert mice (p = 1e^−18^, signed-rank test, 8 mice, 783 cells). Right, population-level scatter plot comparing spike coincidence between HM (blue) and HT (orange) trials in naive mice (p = 0.01, signed-rank test, 4 mice, 409 cells). (**k**) Real vs. predicted spike coincidence (663 neuron pairs). Predicted bilateral spike coincidence was calculated from the unilateral responses. (**l**) Scatter plot comparing real (Hit trials) and predicted spike coincidence (from unilateral responses) across the population of recorded neurons in expert mice (p=3.9e^−76^, signed-rank test, 8 mice, 783 cells). All error bars represent the mean ± s.e.m.

Next, we examined how much of this IH synchrony could be attributed to bottom-up signaling, given the observed changes in BWS and IH touch rate (Fig. 1h, Supp. Fig. 5a, b). To examine the impact of touch kinematics, we calculated IH firing rate as a function of IH touch rate. In naïve mice, IH firing rates weakly increased with increasing IH touch rate, indicating that more synchronous touches evoked marginal gains in IH spike synchrony (Fig. 3d). During Hit trials in expert mice, IH firing rate rapidly increased as a function of IH touch rate (Fig. 3d). Yet, on CR trials, IH firing rates scaled much more slowly, and never obtained a level equivalent to Hits. Naïve mice displayed overall weaker IH firing rates that were nearly equivalent between HT and HM conditions (Fig. 3e, f). Therefore, IH spike synchrony was largely controlled by behavioral relevance.

Next, we sought to determine if touch-evoked spiking was the primary cause of IH synchrony, or if spikes occurring outside the touch period also contributed. To test this possibility, we calculated IH spike coincidence relative to touch onset. To minimize variability, we limited our analysis to bilateral touches with an ITI ≤10 ms. Spike coincidence over time was calculated using the diagonal of the joint peri-stimulus time histograms (jPSTHs, Fig. 3g). Spike coincidence was significantly greater during touches associated with reward, as shown in example neuron pairs (Fig. 3h) and populations of neuron pairs from mice in different behavioral contexts (Fig. 3i). We jittered (±75 ms) the jPSTHs, to identify the periods of time when spike coincidence was greater than a shuffled distribution. As shown in the jPSTHs, this excess spike coincidence primarily occurred in a narrow (∼20 ms) window of time after touch onset, revealing that any spikes occurring outside the brief tactile period made little contribution to IH synchrony. In expert mice, spike coincidence was significantly greater during Hit than CR trials, while in naïve mice, the stimulus configuration (HT/HM) had a marginal effect (Fig. 3j). In naïve mice, spike coincidence was poorly explained by the ITI, while in expert mice, there was a strong linear relationship (Suppl. Fig. 5c). Therefore, IH synchrony was selectively permissive to tactile information. Lastly, we tested if IH spike coincidence during bilateral discrimination was a unique computation, or if it could be replicated solely from its component unilateral touch responses. To do so, we created a predicted jPSTH by shifting the temporal alignment between the unilateral touch PSTHs, so they matched their corresponding sample of bilateral touches (Suppl. Fig. 5d, e). Spike coincidence during bilateral touch was significantly greater than predicted from the unilateral touch responses (Fig. 3k, l & Suppl. Fig. 5e). Taken together, these data reveal that an internal mechanism unlocks IH communication to bind tactile features and drive behavior. Goal-directed changes in spike synchrony were also observed among populations of neurons in the same hemisphere, yet to a lesser degree (Suppl. Fig. 7 a – d).

### Interhemispheric phase-locking is locally enhanced between the superficial cortical layers

To determine if IH communication in S1 was spatially specific or broadly distributed across somatotopic space, we calculated the pairwise phase consistency (PPC) between spikes in one hemisphere and the local field potential (LFP) in the opposite hemisphere (Fig. 4a). Using a laminar 3-shank electrode, we identified the location of the principal barrel column of each whisker based on the amplitude of the touch-triggered LFP during single-whisker stimulation (Fig. 4b). For each neuron that displayed a significant response to the bilateral stimulus, we calculated its phase-locking with the LFP in the opposite principal and adjacent (500 µm horizontal distance) columns (Fig. 4c). In expert mice, we discovered that IH phase-locking was greater during Hit than CR trials, while in naïve mice, IH phase-locking was diminished with no clear stimulus preference (Fig. 4d). The strongest IH phase-locking occurred in the frequency range of mouse whisking (8 – 22 Hz). Across the population of neurons, IH phase-locking was significantly greater during Hit than CR trials, indicating that IH communication was controlled by behavioral relevance (Fig. 4e). Putative pyramidal cells (regular-spiking) displayed significantly greater IH phase-locking than putative fast-spiking (FS) interneurons (Suppl. Fig. 6a), suggesting that FS cells primarily reflected local network dynamics. Across cortical depth, IH phase-locking was strongest between the superficial layers (Fig. 4f & Suppl. Fig. 6b, c). In naïve mice, phase-locking was largely unaffected by cortical depth. Across somatotopic space, phase-locking was significantly stronger with the principal than the adjacent LFP, indicating that phase-locking was spatially specific (Fig. 4g, h). Moreover, we observed a significantly greater decline in phase-locking across somatotopic space during Hit than CR trials, indicating that IH communication was spatially sharpened for the reward-associated stimulus (Fig. 4i). Goal-directed changes in spike-field coupling were also observed within the same hemisphere, yet to a lesser degree (Supp. Fig. 7 e-h). Therefore, IH communication between the superficial layers of the principal barrel columns was focally enhanced during bilateral integration of reward-associated stimuli.

**Figure 4.**
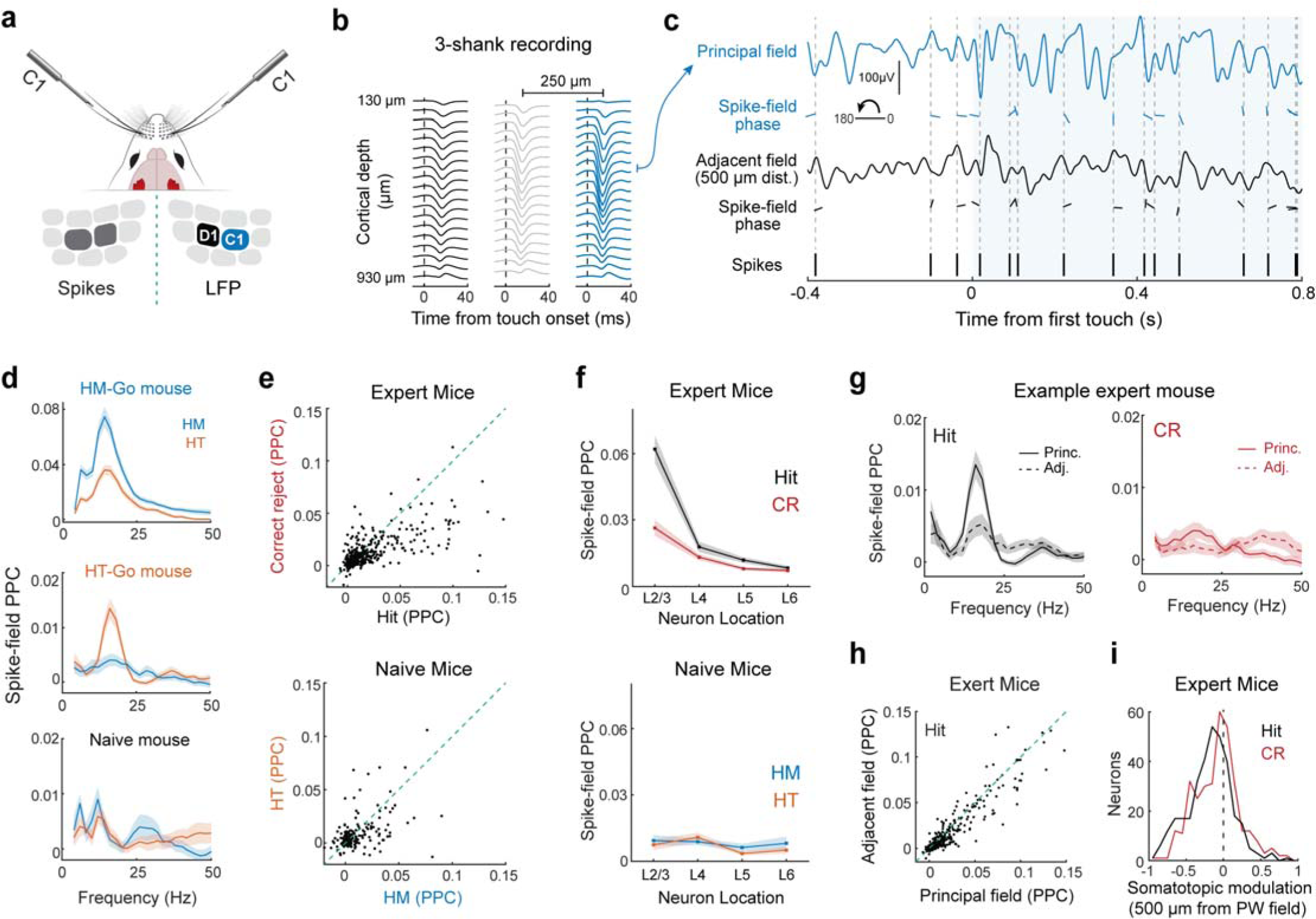
Interhemispheric spike-field coupling is focally enhanced in the superficial cortical layers. (**a**) Schematic of a mouse illustrating the location of spikes in one hemisphere relative to field potentials in the opposite hemisphere during bilateral touch. (**b**) The principal and adjacent barrel columns were identified based on the size of the touch-triggered local field potential (LFP) across our 3-shank laminar recording. (**c**) Example trial showing a neuron spike train from one hemisphere and its corresponding phase relationship to the LFPs in the opposite hemisphere. (**d**) Interhemispheric (IH) spike-field pairwise phase consistency (PPC) in an example HM-Go mouse (top;104 neurons), HT-Go mouse (middle; 63 neurons) and naïve mouse (bottom; 70 neurons). (**e**) Expert and naïve mice population-level IH spike-field PPCs for regular-spiking cells (Expert: p = 7.6e^−15^, signed-rank test, 7 mice, 455 neurons; Naïve: p = 0.08, signed-rank test, 4 mice, 287 neurons). (**f**) PPC according to laminar location of neuron paired with LFP in supragranular layer of opposite hemisphere (Expert mice: 41 neurons in Layer 2/3, 103 in Layer 4, 205 in Layer 5, 106 in Layer 6; Naïve Mice: 55 neurons in Layer 2/3, 84 in Layer 4, 118 in Layer 5, 30 in Layer 6). (**g**) IH spike-field PPCs from one mouse calculated using the LFP that is principal or adjacent (500 µm distant) to the stimulus (47 RS neurons). (**h**) Spatially specific IH phase-locking. A scatter plot comparing the IH phase-locking of neurons to the principal and adjacent (500 µm distant) LFPs (p = 1.25e^−21^, signed-rank test, 7 mice, 455 neurons). (**i**) Spatial sharpening of IH phase-locking (Adj – Princ.). Histogram of the difference in IH phase-locking with the principal and adjacent LFPs (Hit vs. CR, p = 6.45e^−4^, signed-rank test, 7 mice, 333 neurons). All error bars represent the mean ± s.e.m.

### Bilateral integration in S1 neurons reflects task engagement

If an internal mechanism dynamically controls bilateral integration in S1, we hypothesized it would be altered on trials where expert mice incorrectly performed the task. To test this hypothesis, we first analyzed Miss trials, which occurred closer to the end of the experiment (58 ± 5 percentile), when mice were presumably closer to satiety and less engaged. We limited our analysis to mice that performed at least 100 Miss-related touches. Although mice maintained similar whisking and touch kinematics during Miss trials (Fig. 5a, b), the bilateral touch response in a large portion of neurons significantly changed (52%, 250/485 neurons in 5 mice, Fig. 5c – d). The most common change was a decrease in bilateral facilitation (Fig. 5d, e). Next, we focused on False alarm (FA) trials, which occurred closer to the beginning of the experiment (32.5 ± 8.5 percentile), when mice were presumably more eager to receive water reward. We limited our spiking analysis to mice that performed at least 100 FA-related touches. On FA trials, mice whisked with greater BWS, potentially reflecting their expectation of reward (Fig. 5f, 6 mice), yet the strength of active touch was unchanged (Fig. 5g, 3 mice). In the majority of S1 neurons (54%, 186/345 neurons in 3 mice), the bilateral touch response did significantly change, with the most frequent change being an increase in bilateral facilitation (Fig. 5h – j). Taken together, these results reveal that the magnitude of bilateral integration in a large subset of S1 neurons was correlated to task performance.

**Figure 5.**
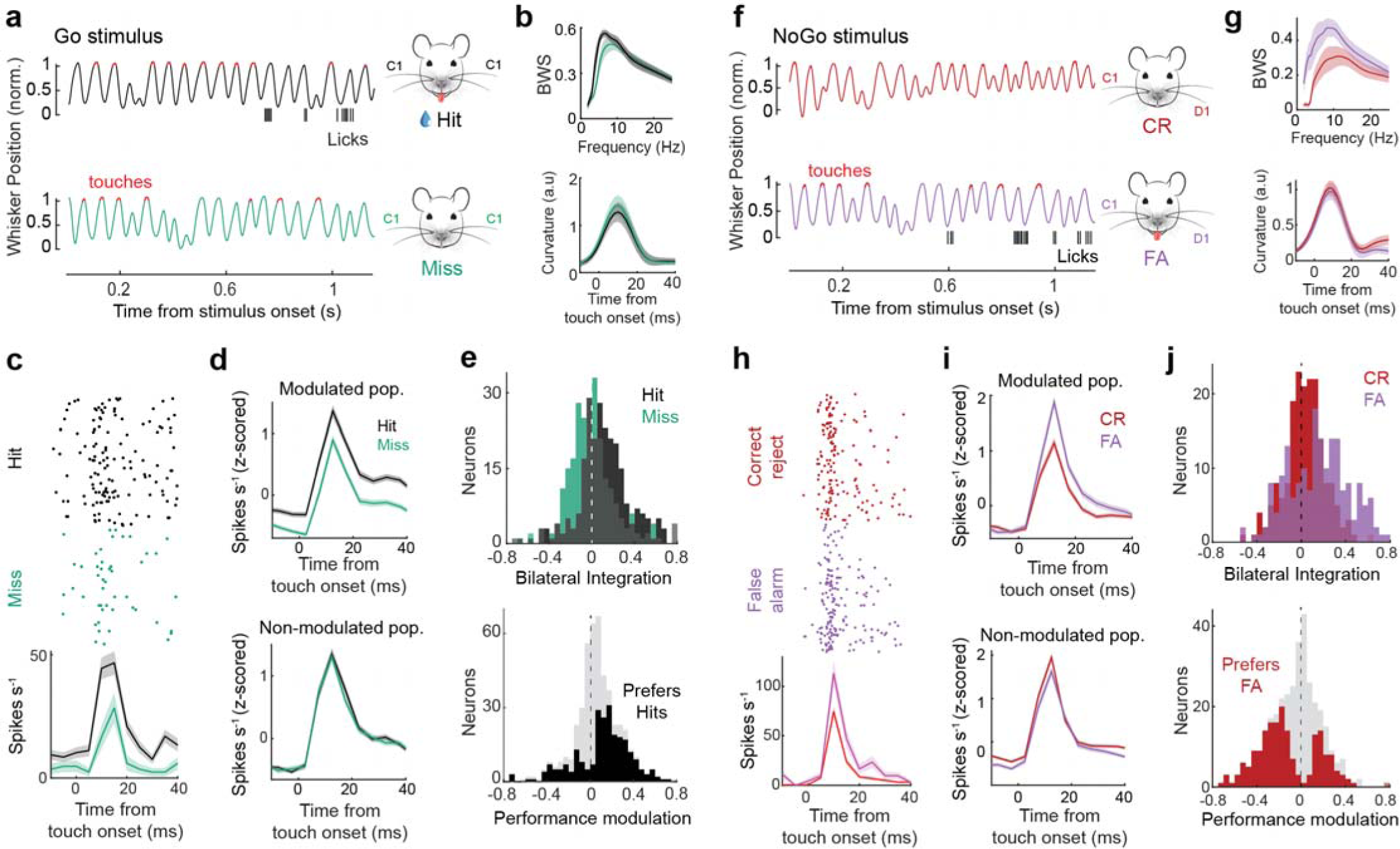
Bilateral integration in S1 neurons is modulated by task performance. (**a**) Example whisker position, touch times, and lick rasters during HM-Go trials that resulted in either a Hit or Miss outcome. Bilateral touch periods are denoted with red patches on the whisker traces. Top: Hit trial; Bottom: Miss trial. (**b**) Top: Average bilateral whisking symmetry (BWS) during hit and miss trials (p = 1e^−4^, paired-sample t-test; 414 samples (9 whisker pairs x 46 frequencies), 6 mice; all whisker pairs with >10 miss trials). Bottom, average whisker curvature for hit and miss trials (p = 0.69; paired-sample t-test; 5 mice). (**c**) Spike raster and PSTHs of firing rate in an example neuron during touches that resulted in a Hit (black) or Miss (green) trial outcome. (**d**) Population averaged touch-triggered firing rates during trials that resulted in a Hit (black) or Miss (green) outcome. Top, neurons that were significantly modulated by Go trial outcome (5 mice, 250/485 neurons). Bottom, neurons that were not modulated by Go trial outcome (5 mice, 235/485 neurons). (**e**) Top, bilateral integration indices of significantly modulated neurons during Hit and Miss trials (p = 2.7e^−10^, paired-sample t-test; 5 mice, 250 neurons). Bottom, performance modulation indices (Hit – Miss) of neurons significantly modulated by trial outcome (black; p = 1e^−08^, one-sample t-test; 5 mice, 250 neurons) and all recorded neurons in these mice (grey; 5 mice, 485 neurons). (**f**) Example whisker position and lick traces during a no-go trial. Top, correct rejection (CR) trial; Bottom, false alarm (FA) trial. (**g**) Top, average BWS during CR and FA trials (p = 5.2e^−32^, paired-sample t-test; 363 samples (8 whisker pairs x 46 frequencies), 6 mice; all whisker pairs with >10 FA trials). Bottom, average whisker curvature at touch during CR and FA trials (p = 0.43; paired-sample t-test; 3 mice). (**h**) Rasters and PSTHs of spiking in an example neuron during touches that resulted in a CR or FA trial outcome. (**i**) Population averaged touch-triggered firing rates during trials that resulted in a CR and FA trial outcome. Top, neurons that were significantly modulated by No-Go trial outcome (3 mice, 186 neurons). Bottom, neurons that were not modulated by No-Go trial outcome (3 mice, 159 neurons). (**j**) Top, bilateral integration indices of neurons significantly modulated by trial outcome (p = 1.4^−8^, paired-sample t-test; 3 mice, 186 neurons). Bottom, performance modulation indices (CR – FA) of neurons significantly modulated by trial outcome (red; p = 2e^−09^, one-sample t-test; 3 mice, 186 neurons) and all neurons in the same mice (grey; 3 mice, 345 neurons). All error bars represent the mean ± s.e.m.

### S1 neurons predict the onset of reward

Recent studies have revealed that activity in the primary sensory cortices is modulated by non-sensory information^46^. We observed a small subset of S1 neurons in expert mice (3.2%, 29/914 neurons) that significantly increased their firing rate immediately preceding reward onset. On Hit trials, these neurons increased their firing rate when the touch period ended, as shown in two example neurons (Fig. 6a, left, black traces). On Miss trials, when the mouse was presumably not expecting reward, firing rates in these same neurons remained at baseline or greatly decreased when the touch period ended (Fig. 6a, left, green traces). These firing rates are in stark contrast to most S1 neurons, which simply returned to their baseline firing rate, irrespective of trial outcome (Fig. 6a, right). The elevated firing rates preceding reward did not occur during touch (Fig. 6b), confirming the non-sensory origin of this activity. Population firing rates demonstrate the pronounced difference in post-touch activity during Hit, Miss, and CR trials (hit/CR: 29 neurons, miss: 23 neurons, Fig. 6c). Most neurons reached their peak firing rates immediately before reward onset (Fig. 6d), while on CR trials, post-touch firing rates were comparable to the pre-stimulus baseline (Fig. 6e). Most reward prediction neurons were in the deeper cortical layers (Fig. 6f), and were putatively excitatory (regular-spiking, 27/29 neurons, Fig. 6g). Taken together, these data reveal a small subset of infragranular S1 neurons that predict reward onset.

**Figure 6.**
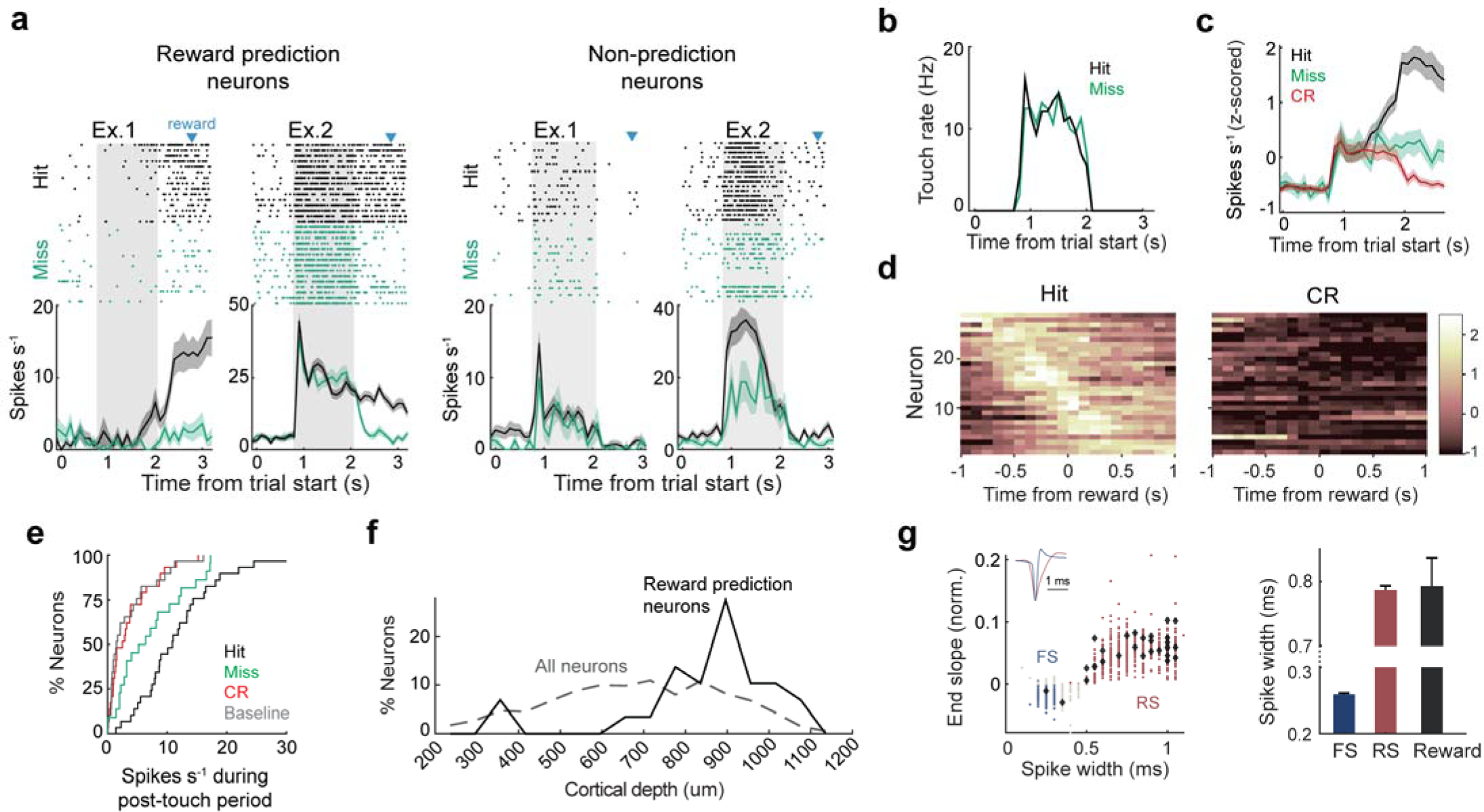
S1 neurons predict reward onset. (**a**) Raster’s and PSTHs of spiking from example reward prediction (left) and non-prediction neurons (right) during Hit and Miss trials in one expert mouse. Gray shadow and blue arrow indicate stimulus presence and time of reward delivery, respectively. (**b**) Average touch rate of the mouse in (**a**) during hit and miss trials. (**c**) Population averaged firing rates (mean ± s.e.m) of reward prediction neurons during hit (black), miss (green), and CR (red) trials (10 mice, 29 neurons). (**d**) Heatmap of normalized activity for reward prediction neurons during hit (left) and CR (right) trials, sorted by the time of maximum activity relative to reward delivery. (**e**) Cumulative percentage of average firing rates for reward neurons during reward anticipation period, defined as the time between stimulus end and reward delivery. (**f**) Distribution of reward prediction neurons (10 mice, 29 neurons; black) and all recorded neurons (10 mice, 914 neurons) across cortical depth. (**g**) Left, scatter plot showing normalized end slope and spike width of individual FS, RS, and reward prediction neurons. Grey dots indicate unclassified neurons. Right, average spike widths of FS, RS, and reward prediction neurons. All error bars represent the mean ± s.e.m.

## Discussion

In this study, we reveal the state- and task-dependent logic of bilateral integration in S1 neurons. Bilateral facilitation of the tactile response emerged during a behavior that required bilateral coordination, and it was lost on Miss trials when expert mice failed to respond to the stimulus. These results challenge and expand upon the longstanding theory of bilateral competition in S1, which is based on neural activity in anesthetized or non-behaving animals. S1 neurons in naïve mice were half as likely to display bilateral facilitation, revealing the importance of behavioral context. We also analyzed the temporal pattern of activity between the somatosensory cortices. In expert mice, tactile evoked spikes became highly synchronized across the hemispheres. In addition, phase-locking between putative pyramidal cells and the opposing LFP was focally enhanced between the superficial layers of the principal barrel columns. This IH spike-spike and spike-field coupling was strongest for stimuli associated with reward and was absent in stimulus-matched naïve animals, indicating that an internal mechanism unlocks IH communication to drive behavior. Taken together, these data suggest that the somatosensory cortices switch from two lateralized spatial representations in a naïve state to a single unified representation during goal-directed bilateral coordination.

In humans, bilaterally symmetrical movements require the posterior (sensorimotor) section of the corpus callosum ^1,3,4^. Movements guided by sensory feedback are enhanced by the expectation of reward ^47,48^. When mice performed our task, they moved their whiskers with enhanced bilateral symmetry during reward expectation. Therefore, their active sensing strategy directly supports a hypothesis of increased callosal communication during our task. Mice deliberately decreased their whisking symmetry after rejecting the bilateral stimulus, further supporting this behavior as a marker of internally motivated bilateral coordination. Since expert mice displayed greater whisking symmetry than naïve mice during trial initiation (pre-stimulus), movement symmetry may be a general strategy for optimizing bilateral sensing ^2,5^. Importantly, rodent whisking is not always symmetrical, and becomes overtly asymmetrical when preparing for an upcoming turn of the head or limbs ^49,50^. The control of whisking symmetry involves the cerebellum, indicating that the synchronization of spikes across S1s and their coupling to the olivocerebellar system could be important for coordinating movement ^51^. Taken together, these results suggest that movement symmetry and its underlying callosal circuits support the neural computations underlying bilateral perception.

In contrast to current literature, we found that contralateral touch responses in S1 are augmented by the addition of ipsilateral touch. By recording from mice across different behavioral contexts, we discovered that bilateral integration is controlled by behavioral relevance. In agreement with this task-dependent model of bilateral integration, coherence between the parietal cortices has been shown to increase during bilateral coordination but decrease during bimanually segregated object manipulations ^36^. If bilateral integration in S1 is truly a flexible process, then it should dynamically change in accordance with task engagement. We tested this hypothesis by analyzing bilateral integration on trials where expert mice made an incorrect choice. On Miss trials, when mice were presumably disengaged, we observed a significant decrease in bilateral facilitation. On False Alarm trials, when mice were presumably eager to receive reward, we observed a significant increase in bilateral facilitation. Contextual changes to S1 sensory responses have been previously observed ^52–54^. The circuit mechanisms supporting such modulations could originate from multiple cortical or midbrain areas that alter the excitability of local circuits in a context-dependent manner ^20,55–62^. For our task, one possibility is that behavioral context controls callosal-mediated inhibition onto the apical dendrites of layer 5 pyramidal cells ^32^, which displayed the highest propensity of bilateral facilitation. Overall, these data reveal how active engagement in bilateral coordination controls bilateral facilitation in S1.

Bilateral perception is thought to emerge from coordinated patterns of activity spread across both cerebral hemispheres. Evidence supporting this hypothesis has been found in the visual cortex, under anesthesia and during behavior ^63–65^. The task-independence of IH synchrony in visual but not somatosensory cortex is potentially explained by inherent differences between vision and somatosensation. Vision is a binocular process, where a central stimulus simultaneously activates both visual hemispheres in cooperation with the corpus callosum ^66^. Conversely, touch is inherently flexible, where the choice between unilateral and bilateral integration is deliberate and strategic. Therefore, the ability to voluntarily switch between stimulus competition and coordination is an important feature of somatosensation. We discovered strong spike-spike and spike-field coupling between the somatosensory cortices, but only in mice that were deliberately sharing information between their hemispheres. IH spike synchrony was controlled by behavioral relevance and was localized to the brief post-touch window, indicating that an internal mechanism unlocks IH communication to bind tactile features. IH phase-locking between putative pyramidal cells and the opposing LFP was focally amplified between the superficial layers of the principal barrel columns, supporting a cell-type and layer-specific mechanism. Taken together, our results establish a novel framework for bilateral coordination in S1, whereby IH networks bind somatotopic features to facilitate goal-directed action.

Collectively, this study provides novel insight into the mechanisms of bilateral integration in S1 and challenges the longstanding model of stimulus competition. The foundation for our work is an innovative behavior where uninstructed movements provide a readout for task engagement and reward expectation. Principally, we discovered that active engagement in bilateral coordination causes the somatosensory cortices to switch from two lateralized circuits into a single unified representation. Future work will focus on dissecting the many circuits and cell-types controlling the state-dependent logic underlying bilateral perception.

## Methods

### Experimental models and subject details

Adult CD-1 mice of both sexes, between 9 to 16 weeks of age were used for all experiments. Mice were group-housed with maximum of 5 per cage and were maintained in reverse light-dark cycle (12 h.-12 h.). All experiments were conducted during their subjective night. All experiment procedures were approved by the Purdue Institutional Animal Care and Use Committee (IACUC, 1801001676A004) and the Laboratory Animal Program (LAP).

### Headplate implantation

Prior to behavioral training, mice were implanted with a custom stainless steel headplate that enabled electrophysiological access to both somatosensory cortices. Mice were anesthetized with isofluorane (2 – 5%) throughout the surgery, while monitoring their respiratory rate and response to toe-pinches, to ensure adequate depth of anesthesia. First, the skin and fur around the head area were disinfected with 70% ethanol, followed by Betadine. The dorsal surface of the skull was exposed using sterilized surgical instruments. The skull and wound margins were covered with a tissue adhesive (3M Vetbond) and the headplate was attached to the skull using dental cement (Metabond). Mice were subcutaneously injected with buprenorphine (0.1 mg/kg) as an analgesic. Mice were given two days of rest before entering behavioral training.

### Behavioral procedure and apparatus

Two days after headplate implantation, mice were habituated to head-fixation on a circular treadmill for one hour per day. After habituation (4 – 7 days), we trimmed all their whiskers except for the four that were used for the task. Mice were initially trained to perform bilateral discrimination using an automated system. The training system contained a circular treadmill, four pneumatically controlled touch surfaces (SMC pneumatics), a lick port, and a headpost for fixation fit inside a light-tight box with white noise. Pistons were manually adjusted to ensure each touch surface entered the movement field of only one whisker. Water was gravity fed to the mouse by opening a solenoid valve (Neptune Research & Development, Inc.). The mice initiated each trial by locomoting on the treadmill for a short distance. A rotary encoder (US Digital) and a piezo sensor (Micromechatronics) attached to the lickport were used to monitor locomotion and licking, respectively.

After habituation to the treadmill, mice were placed on water-restriction and their weight and water intake were closely monitored to ensure they received 1.2 mL of water daily and maintained >80% of their original body weight. If a mouse did not receive the full 1.2 mL of water during the training period, they were given *ad libitum* access to the remaining volume.

During classical conditioning, stimuli were presented (either two HM stimuli for HM-Go or two HT stimuli for HT-Go conditioning) in conjunction with water reward, which was delivered within a randomly selected period 350 – 1150 ms after stimulus onset. After mice learned the association between stimulus presentation and water delivery (as evidenced by anticipatory licking), all four bilateral stimuli were presented, but water was only paired with either the HM or HT stimuli, following their pre-determined designation. Once mice showed consistent anticipatory lick responses for their designated Go stimuli, operant conditioning began. In this stage, mice were required to respond within 1350 ms from stimulus onset to receive water reward. Operant conditioning persisted for 3 – 14 days, until mice learned to withhold licking to the NoGo stimuli. If a mouse made a false alarm (lick response to a No-Go stimulus), it was then required to locomote at least twice the normal distance to initiate the next trial. Mice obtaining a behavioral d-prime ≥1.7 for three consecutive days were considered experts and then moved to behavioral testing on the electrophysiology rig (average d-prime at the end of training = 2.91).

### Reward expectation without bilateral discrimination

A similar behavioral apparatus and training procedure was used to train a separate cohort of mice to associate touch with reward but without performing bilateral discrimination. Five mice were trained to discriminate between unilateral C1 and D1 stimuli. During classical conditioning, a single whisker stimulus (C1) on the right side of their face was paired with water reward. Once mice learned the association between C1 touch and reward (as evident in anticipatory licking), they were moved to operant conditioning and trained to withhold licking during the presentation of the adjacent (No-Go) whisker stimulus (D1). Mice were required to lick for the Go stimulus within 1350 ms from stimulus onset to receive water. Obtaining d-prime ≥ 1.7 for three consecutive days was required to be considered expert and transitioned to the electrophysiology rig. During the electrophysiological recording, unilaterally discriminating mice were periodically presented bilateral stimuli that contained the rewarded C1 whisker. Mice licked in response to this stimulus in expectation of reward. Two additional mice were classically conditioned to associate all bilateral stimuli with reward. All four bilateral stimuli (two HM and two HT) were presented in conjunction with water reinforcement. Once the mice showed anticipatory lick responses for all four bilateral stimuli for three consecutive days, it was considered to have formed a sensory association and moved to operant conditioning.

### Intrinsic Signal Optical Imaging

A day prior to electrophysiology recording, we performed intrinsic signal imaging to locate the C1 and D1 barrel columns on each hemisphere. During the procedure, mice were lightly anesthetized with 1% isoflurane and xylazine (0.3 mg/kg). The skull over the estimated barrel cortex was thinned with a dental drill, and an image of superficial vasculature underneath the transparent skull was obtained with a CCD camera (Retiga R1™, QIMAGING) illuminated by green LED light. Changes in blood oxygen were imaged under red LED illumination while piezoelectric strips wiggled the C1/D1 whisker at 20 Hz for 4 seconds. The site of biggest change in fluorescence was marked as the corresponding barrel and was then registered with the superficial vasculature to guide our electrode penetrations. To protect the brain after imaging, the skull was first covered with a soft silicone gel (Dowsil) and then sealed with a hard-setting silicone (Kwik-Cast™, World Precision Instruments).

### In-vivo electrophysiology

Prior to neural recording, head-fixed mice were habituated to performing the task in the experimental rig for 2 – 3 days. On the day of recording, a 3-shank custom probe (Neuronexus) of 128 channels was lowered into the C1 and D1 barrels in each S1 hemisphere. A micromanipulator (NewScale) lowered the probes into the brain at 75 µm/min. Neural data and experimental signals were acquired at 20 kHz using an RHD 512 recording controller (Intan).

The trial structure was controlled and NI-DAQ toolbox (Matlab) and reward delivery was controlled by an Arduino using custom codes. Neural activity, video frames, and behavioral data during the experiment were synchronized by NI-DAQ triggers sent to the Intan controller.

### Whisker tracking & kinematics

Two high-speed (500 fps) infrared cameras (Photon focus DR1) recorded whisker motion. For post processing, DeepLabCut was used to extract whisker kinematics and obtain touch times ^67^. Four labels were evenly spaced to mark each whisker. 200 video frames were manually labelled for training the network. The DLC network was trained for 300-360K iterations, and the final labels were manually checked for accuracy. Whisker position was calculated for each label for both the whiskers on each whisker pad with reference to a user defined point on the whisker pad. The whisker bend was calculated from the three distal labels on each whisker using Menger curvature (*κ*). Whisker touch frames were identified from when the appropriate whisker label entered a region of interest on the touch surface, which was determined by eye.

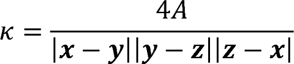

Where ***x, y, z*** are the coordinates of three points on a whisker, and A is the area of the triangle spanned by ***x, y, z***.

### Spike sorting

Recorded neural data were sorted into single units using Kilosort2 and were manually curated using the phy2 GUI (https://github.com/cortex-lab/phy). Spike clusters were classified to be single units based on their cross-correlograms of spike times, location on the electrodes, and waveform characteristics. Only single units were used for all analyses in the paper.

### LFP acquisition

Raw neural data were acquired at 20kHz using an RHD 512 recording controller (Intan). A low pass filter with a cutoff frequency of 300Hz and a notch filter at 60Hz was applied to the raw neural signals. These signals were then down sampled to 500Hz to obtain the final LFPs.

### Discriminability index (d-prime)

Behavioral d-prime (*d*’) is calculated as the difference of z-transforms of hit rate (*Hit*) and false alarm rate (*FA*).

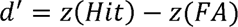

### Bilateral symmetry over time

To quantify the symmetry between bilateral whisker pairs over time (Fig. 1f), whisker positions were lowpass filtered (<30Hz) and the mean whisker angle during the baseline period was subtracted from the entire trial. Then, the difference in position between two opposing whiskers was calculated. Across trials, the absolute value of the difference in whisker positions was calculated and averaged for each stimulus condition.

### Bilateral whisking symmetry (BWS)

To quantify the movement symmetry between bilateral whisker pairs, we calculated their pair-wise phase consistency ^42^. We used custom scripts (Mathworks, 2022) adapted from an LFP-LFP phase synchrony analysis ^68^. Briefly, whisker position data during touch period were first lowpass filtered to reflect whisking frequencies of mice (<30Hz). Then, we applied Morelet Wavelets to the filtered data using MATLAB Wavelet toolbox and obtained a time-frequency representation. Phase angles were calculated using Hilbert transform after datapoints on the same frequency were z-scored across time. Phase synchrony between two whiskers was then quantified using the phase lag between two whiskers with the following definition:

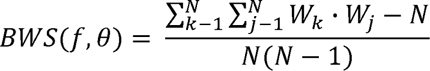

W denotes the phase lag. f and *θ* represent frequency and point in whisking cycle. N is the total number of whisking cycles. Sample sizes for significance testing were calculated as the number of whisker pairs multiplied by the number of frequencies that BWS was extracted from.

### Modulation indices

The contralateral whisker that evoked the largest significant difference in firing rate during post-touch (5 to 35 ms from touch onset) and pre-touch periods (−35 to −5 ms from touch onset) in single-whisker trials was selected as the principal whisker (two-sided Wilcoxon rank-sum test; α = 0.05). HM and HT bilateral response comparisons in single neurons always contained the same contralateral stimulus. Neurons without a significant touch-evoked response were not used in any analysis. Touches were considered bilateral when the IH inter-touch interval (ITI) was less than 26ms, and PW touch time was regarded as onset of bilateral touch. The bilateral integration index was defined as:

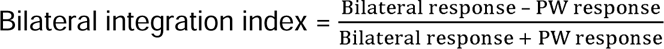

Stimulus preference was calculated by direct comparison between firing rates evoked by two different bilateral touches involving the principal whisker.

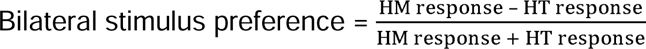

Performance modulation was measured by comparing touch-evoked firing rates during correct and incorrect trials while maintaining the same stimulus.

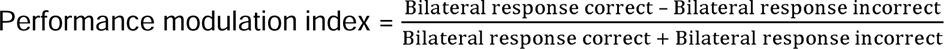

### Classification of best bilateral stimulus

The bilateral stimulus with higher bilateral integration was designated as the best bilateral stimulus of the neuron. If both bilateral stimuli had a bilateral integration index below 0, then the neuron was considered to best respond to the PW stimulus.

### Classification of ipsilaterally responsive neurons

To classify ipsilaterally responsive neurons, we used trials with unilateral whisker contact on the ipsilateral side of the face. Firing rates before and after every ipsilateral touch (−35 to −5ms pre touch, 5 to 35ms post touch) were calculated. We performed a two-sided Wilcoxon signed-rank test on the pre- and post-touch firing rates to determine whether the post-touch response was significantly different from pre-touch period. Neurons with a p-value < 0.01 were classified as ipsilaterally responsive.

### Regular vs. fast-spiking classification

Neurons were identified as regular spiking (RS), or fast spiking (FS) based on properties of their spike waveforms at the electrode site with largest amplitude. Spike width (Trough-to-peak duration), end slope (slope at 0.5ms post trough), and time for repolarization were used as selection criteria ^69–71^. We then performed k-means clustering with these waveform properties to separate RS and FS neurons. Neurons clustered as FS were further checked for their trough-to-peak duration and were removed from the cluster if their spike width was longer than 0.4ms.

### Classification of behaviorally modulated neurons

To classify behaviorally modulated neurons, firing rates after every bilateral touch in correct and incorrect trials were calculated using a 30ms window (5-35ms from touch onset). We performed a t-test on the firing rates comparing responses during correct and incorrect trials to determine the significance of firing rate difference. Neurons with p < 0.05 were considered behaviorally modulated.

### Classification of reward prediction neurons

Reward prediction neurons were determined by calculating post-touch firing rates occurring before reward onset. First, neurons with their greatest post-touch firing rate occurring on hit trials were selected. Then, an ANOVA (Matlab anovan) was performed on each selected neuron comparing hit, correct reject, and PW firing rates during the post-touch period (Tukey test for multiple comparisons). Neurons with a post-touch firing rate significantly greater on Hit trials than all other conditions were classified as reward predictors.

### Interhemispheric spike rates

To calculate interhemispheric (IH) spike rates, a spike triggered average of IH spike intervals was constructed during bilateral touch trials. To create a shuffled distribution of IH spike intervals, spike times of the leader neuron were shifted by a random number up to ±75 ms. To calculate IH firing rates in terms of percent change, we subtracted the shuffled mean from the real distribution and then divided by the shuffled mean. IH firing rates were only calculated between neurons that both showed a significant response to the stimulus.

For the individual neuron analysis (as shown in scatter plots), we calculated the mean IH spike rate of each neuron with all stimulus-responsive neurons in the opposing hemisphere. The mean IH spike rate for each stimulus condition was determined as the absolute value of the change in IH spike rate occurring with an IH lag between ±75 ms. If a neuron significantly responded to both HM or both HT stimuli, the mean IH spike rate change was used for that condition. Mice which had unclear somatotopic map were excluded from analysis. IH touch rates were calculated in an identical fashion to IH firing rates described above. Touch times were used instead of spike times.

### Joint Peri-stimulus time histograms (jPSTHs)

Joint histograms (jPSTHs) of spike coincidence (spikes^2^/sec^2^) were calculated between neurons during Go and No-Go stimulus epochs. Spike coincidence was calculated 0 – 32 ms from the onset of touch. A shuffled distribution (±75 ms) of firing rates was subtracted from the jPSTH to reveal excess spike coincidence. jPSTHs were only calculated between neurons that had a significant response to the stimulus (two-tailed rank sum, p < 0.05). To minimize the impact of IH touch rate on spike coincidence, all jPSTHs were calculated using IH touch intervals ≤10 ms, except where specified otherwise.

### Spike-field pair-wise phase consistency

Spike-field PPC was calculated using the Fieldtrip open-source Matlab toolbox^72^. PPC2 was used, which computes the dot product of phase vectors from different trials containing the same stimulus. This approach controls for within-trial dependencies. Spike-field PPC in the frequency range of 4 to 25 Hz was averaged for each cell to get the PPC value in the population scatter plots. If a neuron responded to both HM or both HT stimuli, the average PPC for that stimulus category was used to perform the Hit vs CR population comparison. The LFP from either the principal or adjacent (500 µm distance) barrel column was used, as noted in the results. Spike-field PPC was calculated using only the neurons that significantly responded to the stimulus, as determined by a ranksum test (p < 0.05, two-tailed) between the pre- and post-touch firing rates. Only mice which had a clear somatotopic map were used in the analysis, as determined by the touch-triggered LFP across the 3-shank electrode. Somatotopic modulation of the interhemispheric spike-field PPC was calculated by comparing phase-locking to the principal and the adjacent LFPs. Only neurons that had PPCs greater than 1e^−4^ were used to calculate somatotopic modulation.

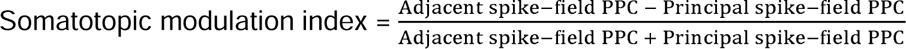

## Data Availability

All data will be made publicly available on Zenodo upon publication. All analysis tools will be made available upon request.

## Acknowledgements

The authors would like to acknowledge the members of the Pluta lab, Daniel Butts, Julia Veit, and Hillel Adesnik for providing valuable feedback on the manuscript.

