## Supplemental figures for "Bilateral integration in somatosensory cortex is controlled by behavioral relevance"

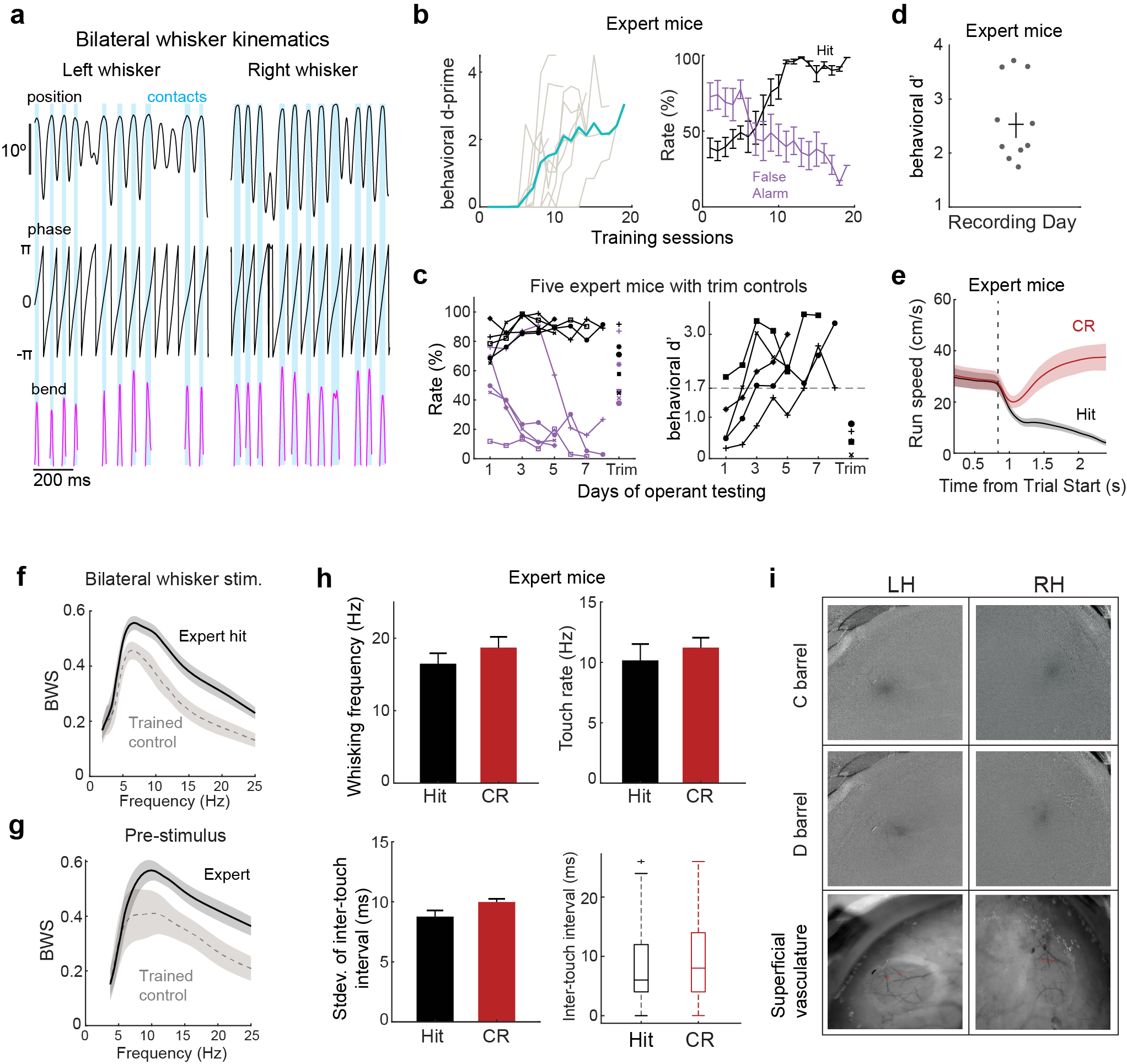

**Supplementary figure 1. Task performance and behavioral dynamics during bilateral discrimination.** (**a**) Example bilateral whisker kinematics. Top: Whisker position. Middle: Whisker phase. Bottom: Whisker bend. Blue shadows indicate touches. (**b**) Left: d-primes of all recorded mice over training sessions (10 mice). Right: Average hit and false alarm rates of the mice over training sessions. (**c**) Average hit and false alarm rates (left) and d-prime (right) of 5 expert mice over days of operant conditioning, followed by a control session with all whiskers trimmed. (**d**) D-prime of expert mice on the recording day (10 mice). (**e**) Average run speed during hit and correct rejection (CR) trials (10 mice). (**f**) Bilateral whisking symmetry (BWS) for all expert mice during hit trials in black and trained control mice in gray (p = 6.3e^-18^, two-sample t-test; 920 samples for hit trials in expert mice (20 whisker pairs in 10 mice) and 552 samples for trained control mice (12 whisker pairs in 5 mice)). Out of five trained control mice, four were performing unilateral whisker discrimination task. Bilateral stimulus trials that included the Go whisker stimulus were used. One mouse was classically trained to expect reward during both HM and HT whisker stimuli. (**g**). BWS of all expert mice (black, 10 mice) and all trained control mice (gray) during the pre-stimulus period (p = 5.3e^-09^, two-sided Wilcoxon rank-sum test; 340 samples from 10 expert mice; 204 samples from 6 trained control mice). (**h**) Whisker kinematics during hit and CR trials (10 mice). From top left to bottom right: average whisking frequency, average touch rate, standard deviation of interhemispheric touch intervals, and box plot of interhemispheric touch intervals. (**i**) Example images from intrinsic imaging on wS1 (top and middle) to locate barrel columns corresponding to whiskers used in the task. The locations of intrinsic signals were registered with images of the superficial vasculature to guide electrode placement (bottom).

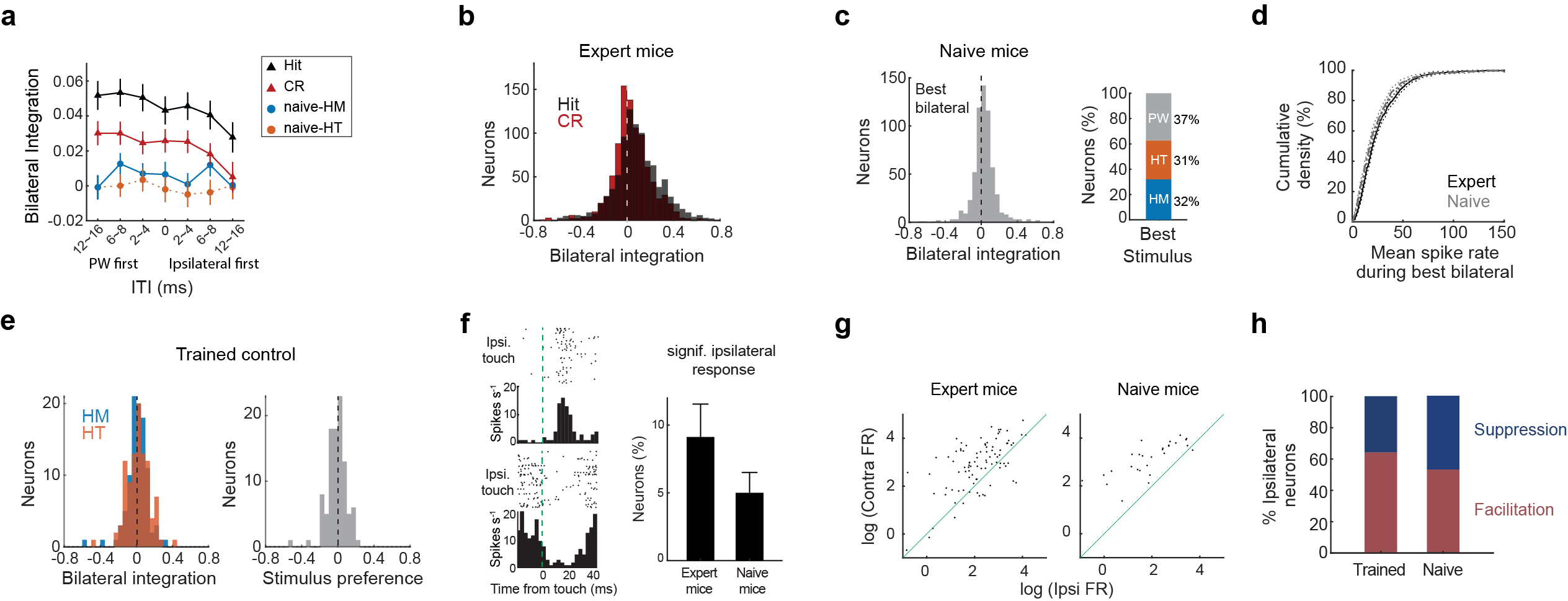

**Supplementary figure 2. Context-dependent bilateral facilitation is maintained across different touch dynamics**. (**a**) Average bilateral integration indices of neurons depending on interhemispheric touch intervals (ITI). Hit and CR are shown for expert mice (10 mice, 914 neurons), and HM and HT for naïve mice (7 mice, 609 neurons). ITI was calculated as difference in time between PW and ipsilateral touch. (**b**) Distributions of bilateral integration indices for hit and CR conditions in expert mice after controlling for numbers of touches included in the calculation (p = 5.7e^-16^, paired-sample t-test; 10 mice, 914 neurons). Index was calculated using the same number of touches within each trial (equal to the mean number of touches in Hit trials). Touches that occurred later in the trial exceeding the mean touch number were excluded. (**c**) Left: Bilateral integration index using best bilateral stimulus for neurons from naïve mice. Right: Percentage of neurons categorized as HM-, HT-, and PW-best. (**d**) Cumulative density of the largest bilateral response in expert mice (10 mice, 914 neurons; black) and in naïve mice (7 mice, 609 neurons; gray). (**e**) Left, bilateral integration indices in mice expecting reward but not performing bilateral discrimination during HM (p = 0.4, one-sample t-test; 3 mice, 99 neurons) and HT touch (p = 0.1, one-sample t-test; 3 mice, 99 neurons). Right, bilateral stimulus (HT vs. HM) preference in mice expecting reward but not performing bilateral discrimination (p = 0.31, one-sample t-test; 3 mice, 99 neurons). (**f**) Left: Histograms of spiking aligned to the onset of unilateral touch with the ipsilateral whisker in two example neurons. Right: Percentage of neurons showing significant response to ipsilateral touches in expert (left) and naïve (right) mice (p = 8.6e^-4^, Fisher’s exact test; expert: 10 mice, 914 neurons; naïve: 7 mice, 609 neurons). (**g**) Scatter plots comparing unilateral responses to ipsilateral and contralateral touch in neurons with a significant ipsilateral response. Left, expert mice (91 neurons; 10 mice). Right, naïve mice (29 neurons; 7 mice). (**h**) Percentage of neurons with significant spike facilitation or suppression during ipsilateral touch. Percentages in trained mice and naïve mice are shown.

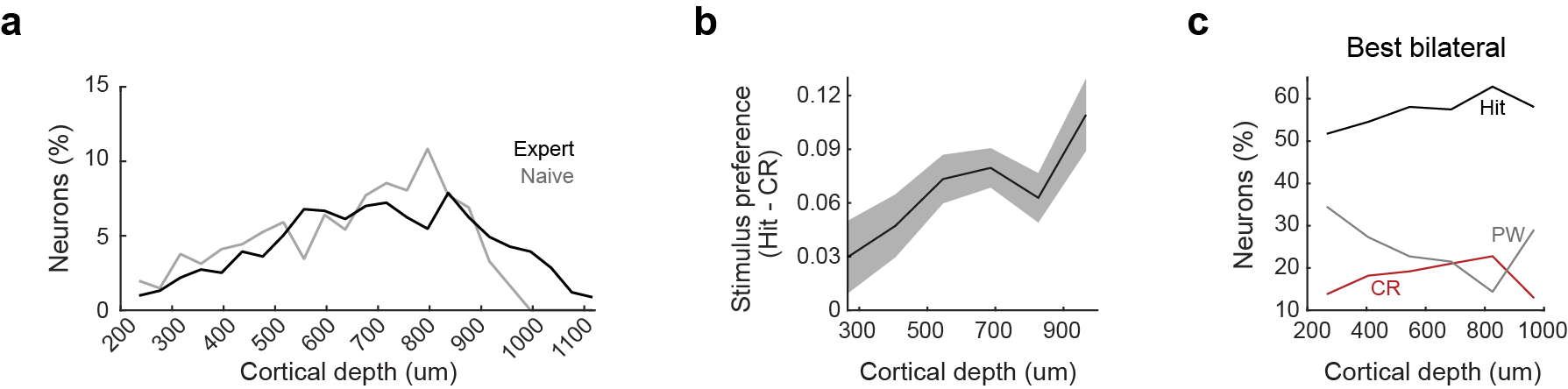

**Supplementary figure 3**. **Bilateral stimulus preference increases with cortical depth**. Characterizing S1 activity across cortical depth. (**a**) Distribution of all recorded neurons across cortical depth in expert mice (10 mice, 914 neurons; black) and in naïve mice (7 mice, 609 neurons; gray). (**b**) Average stimulus preference of neurons in all expert mice across cortical depth (10 mice, 914 neurons). (**c**) Best stimulus of neurons in expert mice across cortical depth (10 mice, 914 neurons).

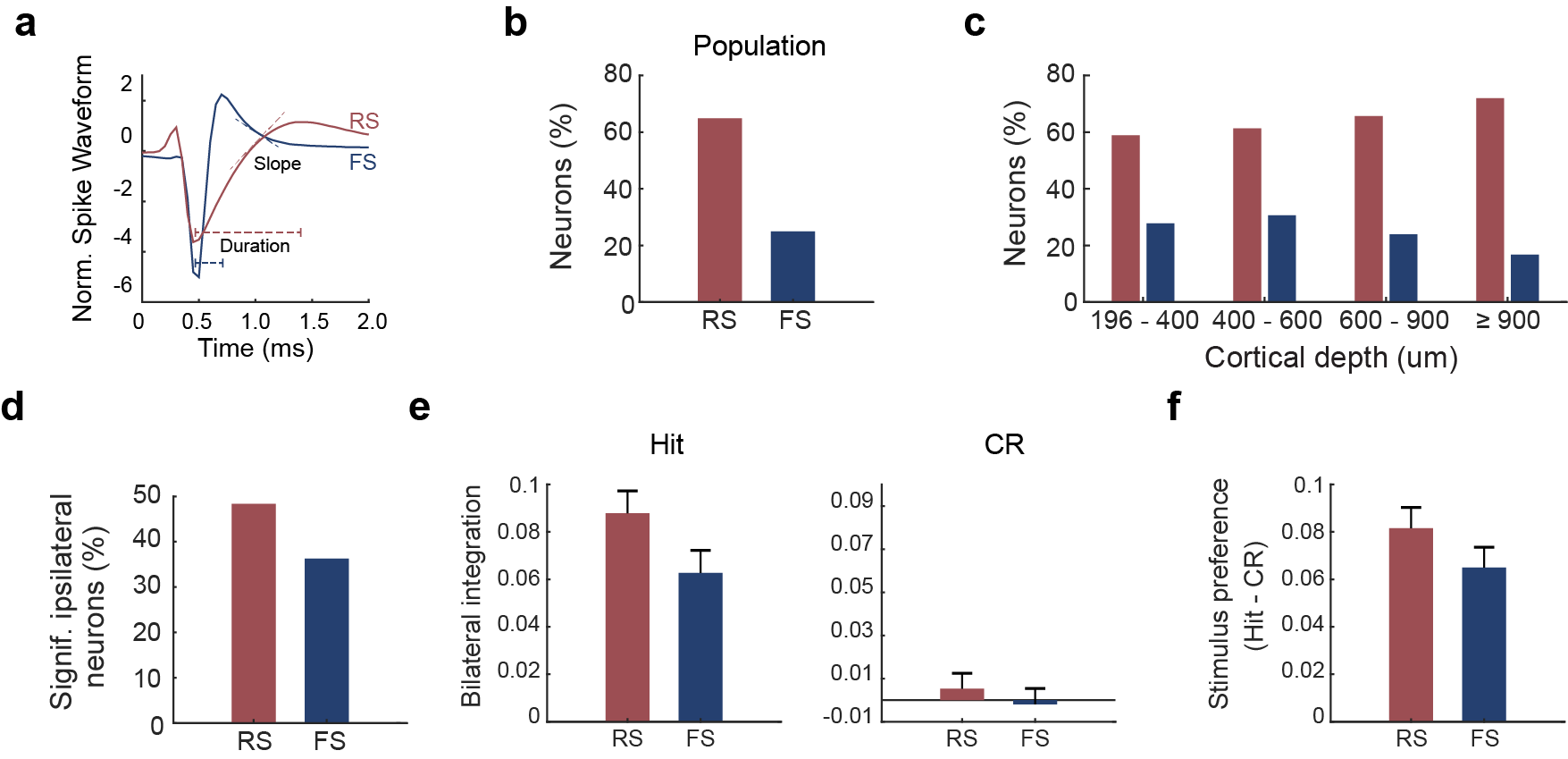

**Supplementary figure 4. Bilateral integration is similar between regular and fast-spiking neurons.** Characterizing neural activity comparing RS and FS neurons. (**a**) Example waveforms of RS and FS neurons, showing 2 of the criteria used for classification. (**b**) Percentage of RS and FS cells in the population of recorded neurons (10 mice, 914 neurons). (**c**) Percentage of RS and FS neurons across cortical depth. (**d**) Percentage of RS and FS cells among significantly ipsilateral neurons. (**e**) Average bilateral integration for RS and FS neurons during hit (left) and CR (right). (**f**) Average stimulus preference index for RS and FS neurons comparing hit vs. CR.

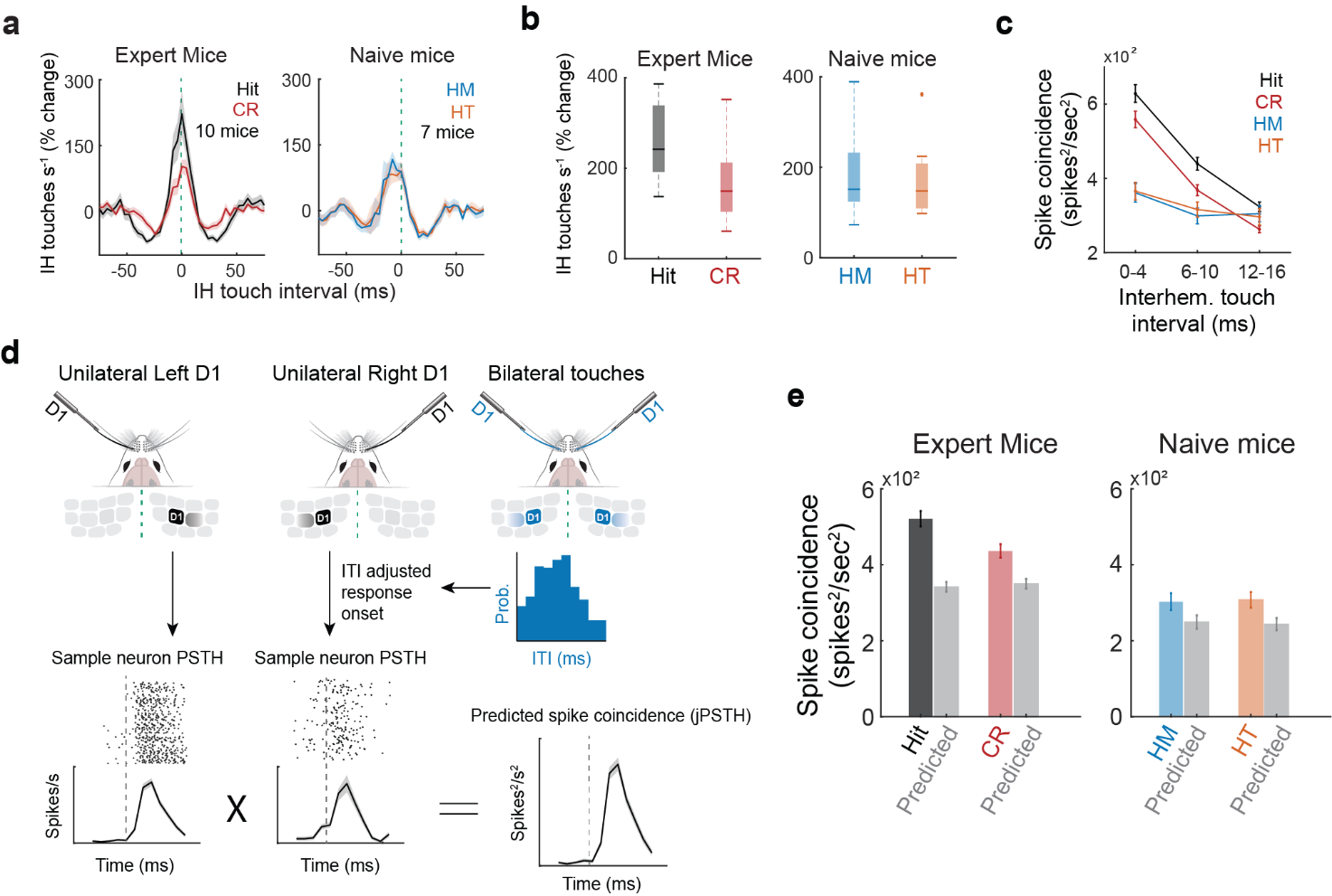

**Supplemental Figure 5. Bottom-up signaling cannot explain the enhanced interhemispheric synchrony in expert mice.** (**a**) Interhemispheric touch rate as a function of touch lags for expert (n=10 mice) and naïve mice (7 mice). (**b**) Box plot of IH touch rates for hit and correct reject trials in expert mice (p=5.2e^-4^, 2-sample t-test, n = 10 mice) and naïve mice (p = 0.76, 2-sample t-test, n = 7 mice). (**c**) Relationship between IH touch interval (ITI) and spike coincidence (jPSTH) in expert and naïve mice. (**d**) Schematic illustrating the generation of predicted spike coincidence from the unilateral touch-triggered PSTHs. The real bilateral ITIs were used to match the temporal relationship of the unilateral touches to the actual bilateral stimulus. (**e**) Bar graphs of average spike coincidence across the conditions (real vs. predicted) in expert (n= 8 mice, m= 783 cells) and naïve mice (n= 4 mice, m=409 cells).

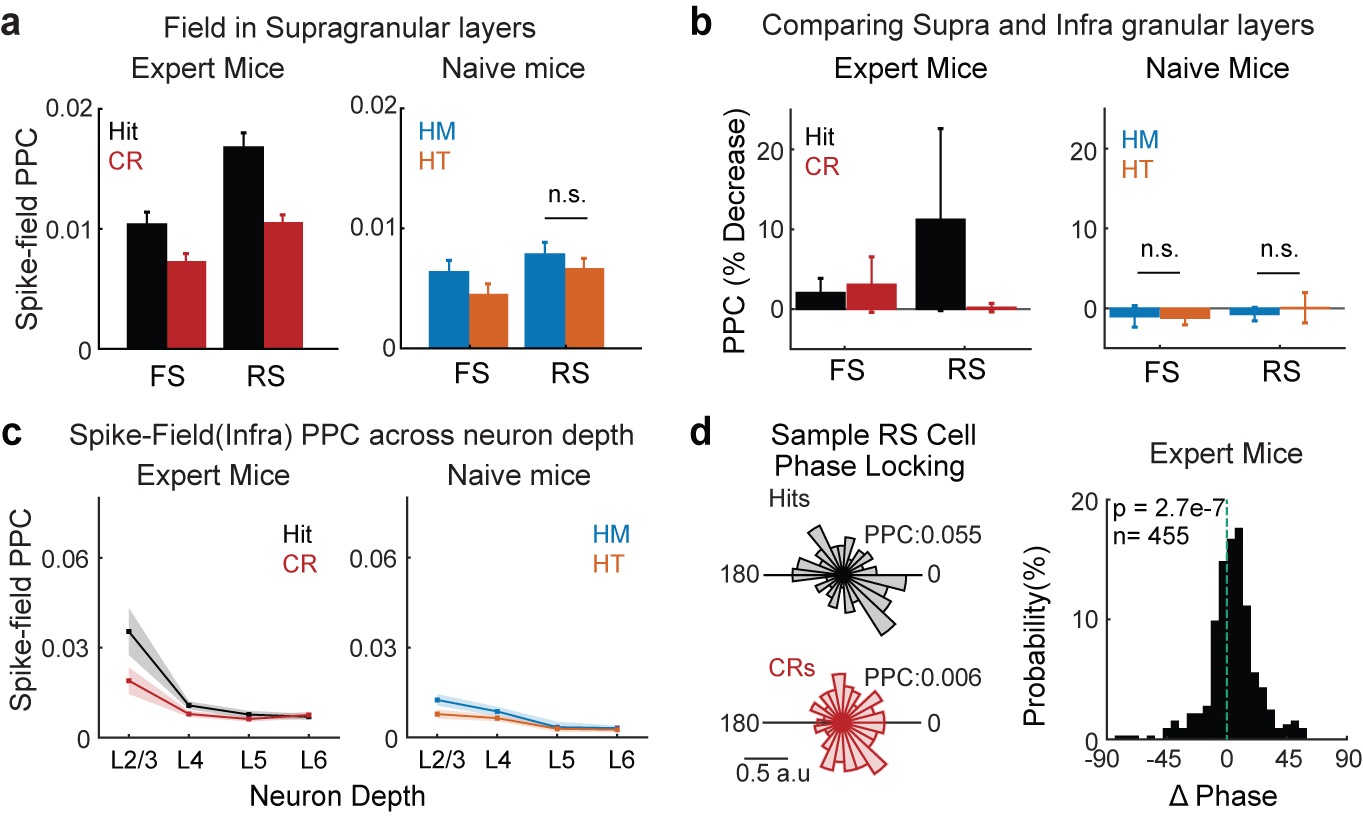

**Supplemental Figure 6. Regular-spiking cells are more strongly coupled to the opposite hemisphere.** (**a**) Interhemispheric (IH) spike-field PPC for fast and regular spiking cells in expert (FS cells: 138, RS cells: 455, 7 mice) and naïve mice (FS cells: 78, RS cells: 287, 4 mice). (**b**) Percent decrease in PPC between supragranular and infragranular field potentials in expert and naïve mice. (**c**) IH spike-field PPC according to laminar location of neuron and using the infragranular field potential. Expert mice: 41 neurons Layer 2/3, 103 in Layer 4, 205 in Layer 5, and 106 in Layer 6. Naïve Mice: 55 neurons in Layer 2/3, 84 in Layer 4, 118 in Layer 5, and 30 in Layer 6. (**d**) Left, histogram of spike-field phase relationships in an example neuron for Hit and CR trials. Right, change in phase preference between Hits and CR in population of RS cells of expert mice (p=2.7e^-7^, signed-rank test, n=455, 7 mice).

**Supplemental Figure 7. Intrahemispheric synchrony is enhanced by goal-directed processing.** (**a**) Left, inter-neuronal firing rates between neurons in the same hemisphere in an example expert mouse. Right, mean area of inter-neuronal synchrony for all expert mice (p=3.7e^-23^, signed-rank test, 867 neurons, 9 mice). (**b**) Same as in (**a**) but for naïve mice (p=0.87, signed-rank test, 744 neurons, 6 mice). (**c**) Left, joint-PSTHs between neurons in same hemisphere of an example expert mice. Right, All responsive neurons in expert mice (p = 2.3e^-36^, signed-rank test, 867 neurons, 9 mice). (**d**) Same as in (**c**) except for in naïve mice (p = 0.87, signed-rank test, 744 neurons, 6 mice). (**e**) Left, within hemisphere spike-field PPC for RS cells in an example expert mouse (n=134), Right, all stimuli responsive RS cells (p = 2.3e^-11^, signed-rank test, 810 neurons, 9 mice). (**f**) Same as in (**e**) except in naïve mice (p = 0.004, signed-rank test, 743 neurons, 6 mice). (**g**) Left, within hemisphere spike-field PPC according to horizontal location of field potential in an expert mouse (98 neurons). Right, all stimuli responsive cells (p = 1e^-11^, signed-rank test, 566 neurons, 9 mice). (**h**) Same as in (**g**) except in naïve mice (p = 4.7e^-8^, signed-rank test, 499 neurons, 6 mice).
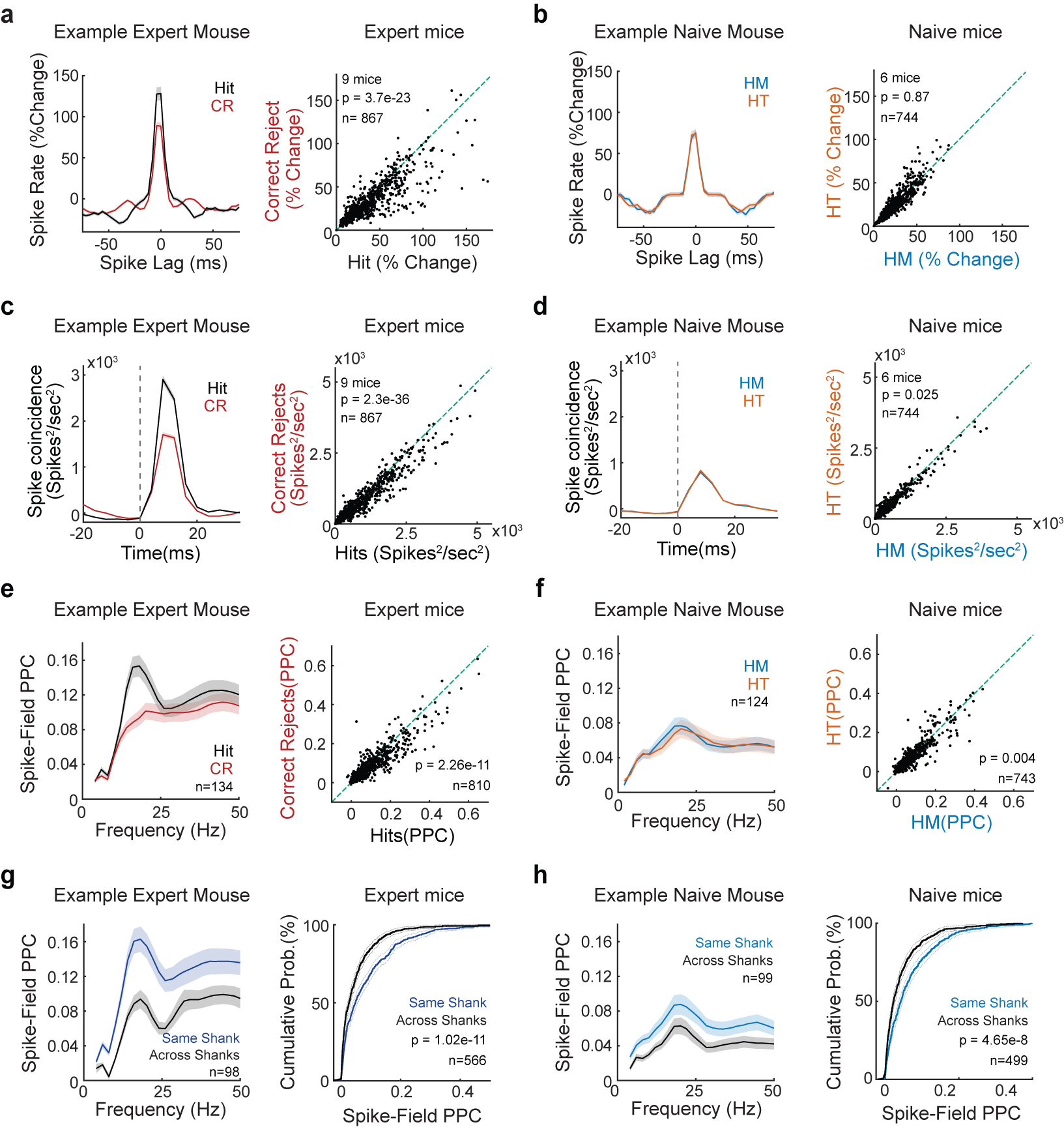
